# ViTAMIn-O: Democratizing computer vision-based machine learning for stem cell research

**DOI:** 10.64898/2026.06.01.726000

**Authors:** Ferhat Hamurcu, Markus Breunig, Arpad Varga, Bianka Bosch, Jessica Lindenmayer, Anjan T. Kanakappady, Kevin Achberger, Natalia Pashkovskaia, Alexander Kleger, Stefan Liebau, Stefanie Klingenstein, Moritz Klingenstein

**Affiliations:** Institute of Neuroanatomy and Developmental Biology (INDB), University Tübingen, Tübingen, Germany; Institute of Molecular Oncology and Stem Cell Biology, Ulm University Hospital, Ulm, Germany; Core Facility Organoids, Ulm University, Ulm, Germany; Division of Interdisciplinary Pancreatology, Department of Internal Medicine I, Ulm University Hospital, Ulm, Germany

**Author notes:** These authors contributed equally (Stefanie Klingenstein, Moritz Klingenstein). Senior author (Stefanie Klingenstein, Moritz Klingenstein).

## Abstract

Deep Learning (DL) holds exciting potential in automating the prediction of organoid differentiation results. Nevertheless, current models lack adaptability, openness, and robustness in performance. Additionally, broad employments of predictive models in wet-lab settings necessitate machine learning expertise, often not readily available in biologically oriented laboratories. To offer an intuitive solution, we present ColabViTAMIn-O, a code-free platform together with ViTAMIn-O. ViTAMIn-O is a fully open organoid-specific DL model trained and tested on a total of 34 organoid categories, incorporating annotated images across transmitted light microscopy (TLM) modalities at single-organoid resolution. It is adaptable to downstream prediction tasks of varying dataset sizes and outperforms established models even with linear-probing. It performs reliably within a few-shot framework and is even extensible to human embryo TLM imaging data at single specimen level. By releasing our platform, centralized model hub, and datasets, we hope to encourage broader deployments of specialized DL models in stem-cell laboratories.

## INTRODUCTION

Organoid technologies encompass a diverse range of in vitro model systems, including those derived from adult human tissue samples and patient-specific tumors^1,2^. Alongside these primary tissue-derived models, pluripotent stem cell-derived systems have emerged as insightful tools to study human development and disease. The discovery of induced pluripotent stem cells (iPSCs)^3,4^ and the efficiency of minimally invasive reprogramming techniques^5–9^, alongside efficient protocols to derive tumor cells from human tissue samples have catalysed the emergence of organoids as highly relevant in vitro model systems. Organoids are self-organizing three-dimensional cell aggregates derived from diverse cell populations, such as iPSCs, which recapitulate key steps in early development^10,11^. However, organoid differentiation is inherently characterized by its probabilistic nature, among other things, due to the difficulty of achieving precise spatio-temporal mimicry of in vivo morphogenetic events^12^. Consequently, the refinement and establishment of new differentiation protocols remain a time and cost intensive pursuit.

The Transformer revolution^13,14^ has enabled Computer Vision (CV) models to accommodate for ever larger datasets in medical contexts^15,16^. In contrast to Convolution Neural Networks (CNNs), transformer-based architectures tend to generalize more effectively over larger quantities in imaging data, substantiating their role as the backbone for a variety of biomedical vision foundation models^17–19^. Diverse and sufficiently large quantities in imaging data have precipitated the rise of predictive CV models across different biomedical fields, from bench^20–23^ to bedside^24–28^. Nevertheless, the sparsity in systemically annotated imaging data remains a central issue in employing transformer-based architectures in stem cell biology. Furthermore, existing large scale organoid imaging datasets disregard morphological intricacies on single organoid level by providing whole well displays or focus on organoid detection^29,30^ and segmentation^31–34^. These limitations further impede subpopulation classification and cell fate prediction. Though several groups have investigated the feasibility of implementing deep learning methods to predict organoid specific cell fate trajectories^35–40^, these models were trained on homogenous, often lab-internal datasets tailored to singular organoid types. Notably, deployment of these highly specialized models remains largely restricted to specialized laboratories, hindering explorative and translatory research implementation.

Taken together, there is a need for a standardized easy to use analysis pipeline that enables early, non-invasive assessment of global organoid viability and fate prediction assessment, offering a scalable, cost efficient and user-friendly platform, which ideally mitigates the need of machine learning or even prior coding experience.

To overcome these constraints and accelerate progress in a variety of stem-cell based optimization efforts in a fully open and user-friendly way, we present ColabViTAMIn-O, hosting ViTAMIn-O (Vision Transformer based Artificial Morphological Intelligence on Organoids) and the ViTAMIn-O Model Hub. ViTAMIn-O is a novel, fully open generalist CV model, easily customizable to new lab-internal datasets through the one click platform ColabViTAMIn-O for stem-cell model classification and differentiation prediction tasks. It was pretrained in a supervised manner making usage of nine diverse organoid imaging datasets, incorporating knowledge from seven carefully selected publicly available datasets and two newly generated lab-internal datasets. The training dataset consists of two human brain^41,42^, hypothalamic-pituitary^35^, intestinal^43^, three colon^44^, pancreatic ductal adenocarcinoma (PDAC), salivary adenoid cystic carcinoma (ACC), distal airway epithelia, murine small intestine (from here on summarized as ACLMP^45^), human anterior placode^46^ and otic^47^ organoids, encompassing a total of n = 32,773 images for the training set n = 24,751, validation set n = 3,611, internal test set n = 4,411 and external test set of n = 1,260 organoids as well as a total 7,844 annotated blastocyst^48^ and embryo images^49^ through multiple developmental stages. ViTAMIn-O builds upon the Swin transformer^50^ architecture and utilizes a teacher-student network^51^ during the pretraining phase. Under this premise, the student was fed extensively augmented images, whereas the teacher was trained using unaltered images, with its weights updated using exponential moving averages (EMA)^52–54^. Through extensive comparative benchmarking experiments, illustrating its extensibility in predictive capabilities to embryo datasets, introducing an explainable and intuitive way of understanding the model’s decision-making process^55^ and making all of this openly accessible through ColabViTAMIn-O and the integrated ViTAMIn-O Model Hub, we aim to catalyze further developments of the next generation of robust and widely deployable computationally guided tools concerning stem-cell technology.

## RESULTS

### Main architecture and platform applications

ViTAMIn-O is a computer vision model trained through a supervised teacher-student network guided pre-training phase, leveraging a Swin transformer model and incorporating a wide-ranging set of organoid imaging data (Figure 1). It was trained by differentiating between 28 organoid classes derived from 11 organoid types by cyclically reiterating through these datasets by an either a) heavy data augmentation pipeline provided to the student model or b) only resized images to the teacher model (Figures 1A and 1D). ViTAMIn-O excels across various supervised organoid prediction tasks, ranging from machine learning algorithms such as logistic regression or k-nearest neighbour to deep learning domains such as few-shot learning, linear-probing or fine-tuning. After benchmarking our model against other state of the art deep learning models (Figure 2), we then moved forward and investigated ViTAMIn-O’s performance against various biological microscopic imaging settings, across never before seen stem cell model systems (Figure 3). We show this by probing our model on three external organoid datasets, two more external embryonic imaging datasets and additionally incorporate an explainable AI module on our platform (Figure 3 and Figure 5). Moreover, we provide with ColabViTAMIn-O an easy-to-use, non-expert computational platform enabling wet-lab researchers to use their individual images to derive new morphologically guided hypotheses on their model systems through Automated Concept-based Explanation^55^ on organoids (OrgACE). We demonstrate its practicability through employing this workflow on an otic organoid model and validate the results through immunofluorescence stainings of key protein markers in otic organoid differentiation and qualitatively match the spatial expression profile of organoids with the predicted OrgACE output (Figures 5C and 5D). Lastly, by integrating the ViTAMIn-O Model Hub within the ColabViTAMIn-O infrastructure, we provide a centralized model repository. This feature enables researchers to upload and share trained models, supporting both independent analyses and collaborative research efforts. Ultimately, we hope to democratize the ability to train and deploy highly capable vision guided AI models for stem cell-based phenotypic screens, omitting the need of expertise in machine learning domains or advanced coding experience in its entirety.

**Figure 1.**
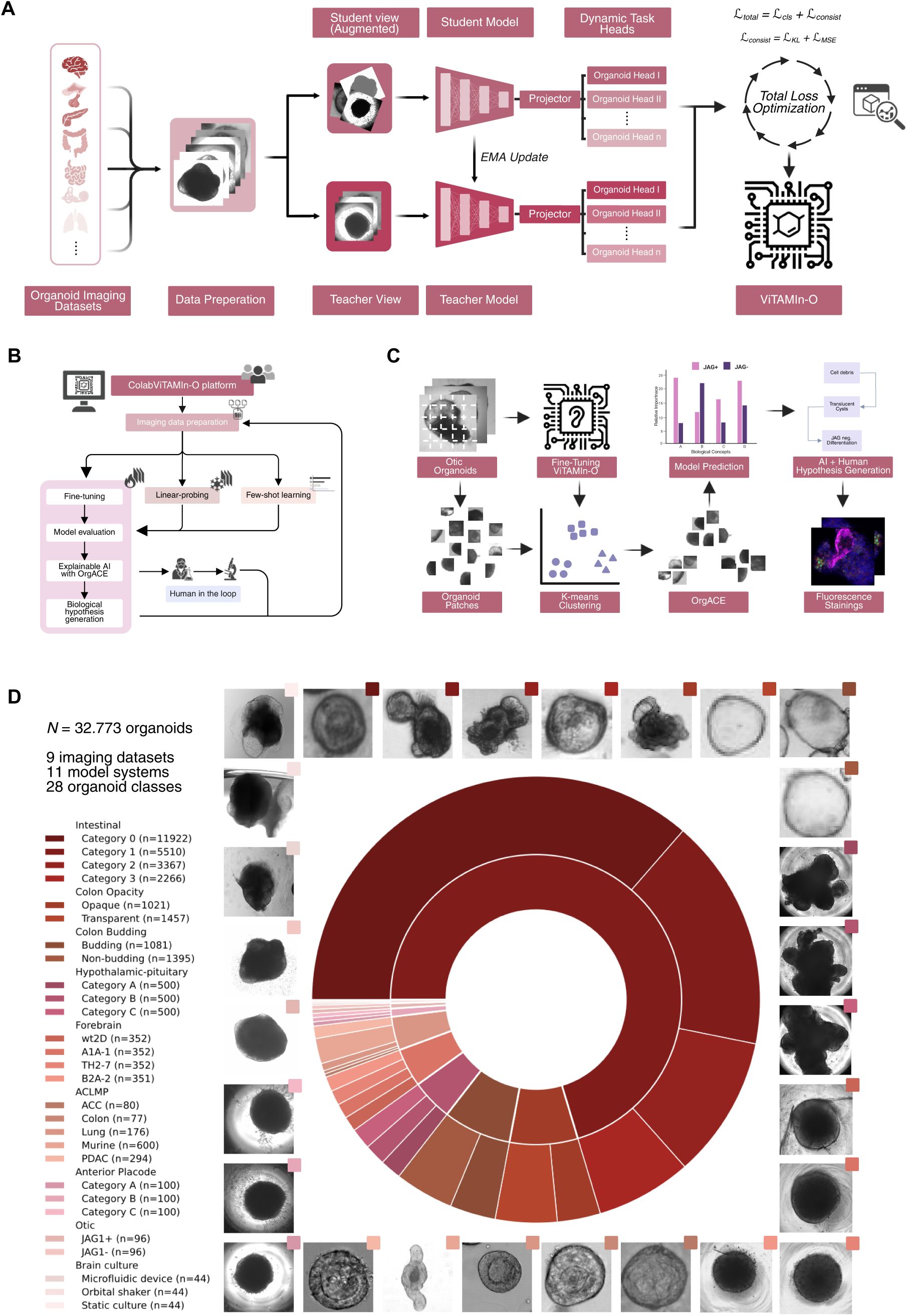
Training of ViTAMIn-O, ColabViTAMIn-O structure and OrgACE. (A) ViTAMIn-O was pretrained by accruing nine heterogenous organoid datasets, incorporating brightfield and phase contrast imaging modalities of organoids in a total of 28 categories. It was trained by utilising a teacher-student network with multitask heads. The student is, contrary to the teacher, trained on heavily augmented training images. After each iteration, the updated weights of the student are transferred to the teacher through exponential moving averages (EMA). (B) ColabViTAMIn-O is an intuitive platform, offering various deep learning applications, hosted on a Google Colab environment. Through a one click approach, users can train individualized models by fine-tuning, linear-probing or few-shot learning, mitigating the need of coding experience in its entirety. These models are easily downloadable and deployable for future differentiation prediction efforts through the ViTAMIn-O Model Hub integration as well as verifiable through Automated Concept-based Explanation on Organoids (OrgACE), offering the possibility of a closed loop cycle of imaging data generation, AI prediction and hypothesis generation of stem cell differentiation variability. (C) Employing ViTAMIn-O and OrgACE to decipher morphological correlates of JAG1 expression in otic organoids. Otic organoids are divided into image patches, representing biologically informed morphological patches and mapped through a K-means clustering algorithm on a fine-tuned ViTAMIn-O model. Enabled by the OrgACE algorithm, biological concepts associated with or without JAG1 protein expression are clustered into distinct regions. Through the ColabViTAMIn-O platform, users can discover these morphological concepts such as the appearance of translucent cyst structures and associate these with spatial protein mapping techniques such as immunofluorescence analysis. (D) Composition of the ViTAMIn-O pretraining dataset. The donut chart illustrates the partitioning of 32,773 single organoid transmitted light microscopy-derived images, encompassing 28 morphological classes across 11 distinct organoid model systems. The inner ring visualizes the relative contribution of overarching global datasets (e.g., intestinal, colon budding), while the outer ring details the specific per-class distributions (e.g., Category 0, Category 1). Absolute quantitative shares for each category are provided on the left.

**Figure 2.**
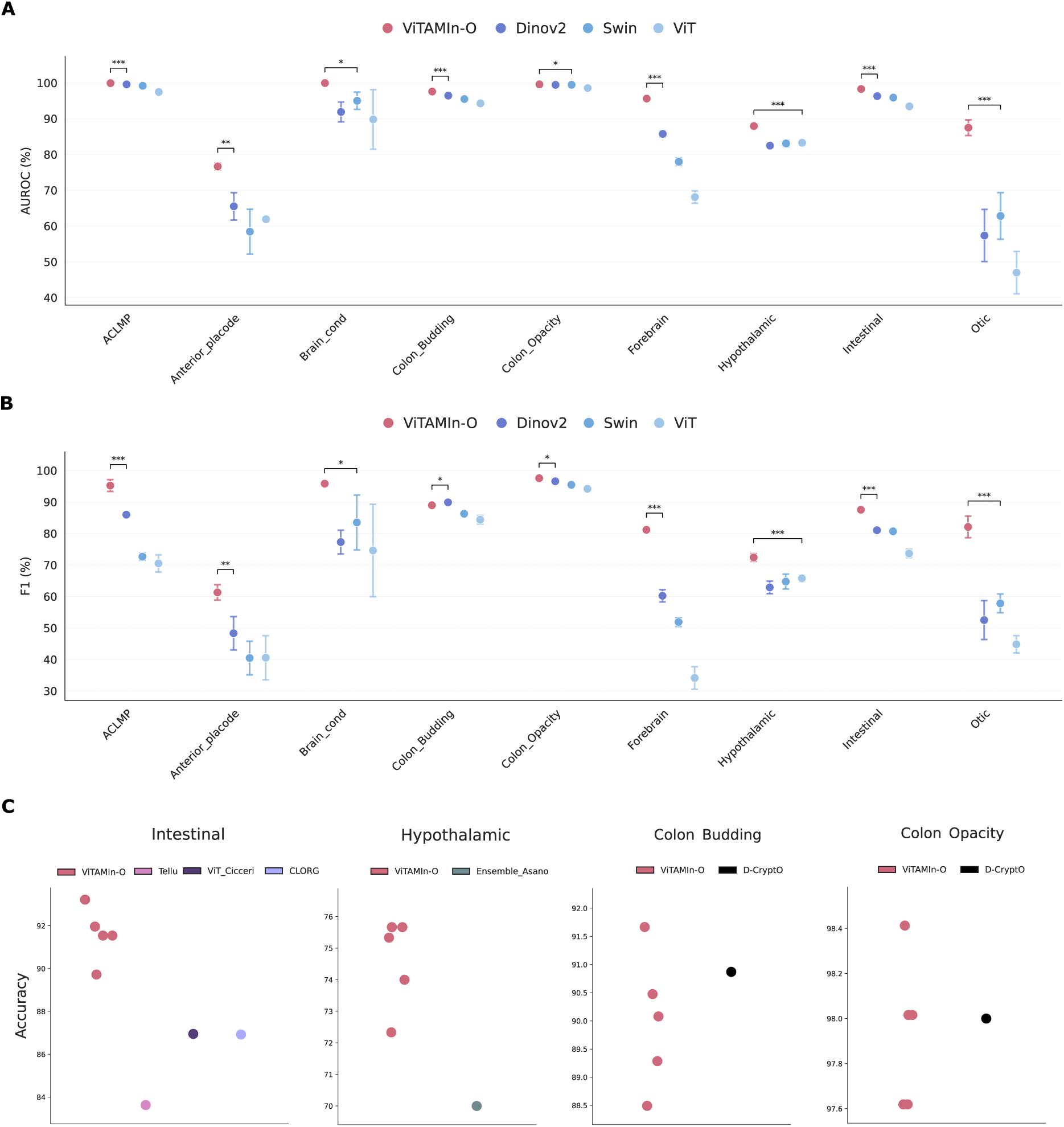
Performance benchmarking of ViTAMIn-O across organoid model systems and learning configurations. (A) AUROC comparisons under linear-probing configuration. ViTAMIn-O outperformed the next best large pretrained models (DINOv2, Swin, or ViT) across all nine global organoid prediction tasks. (B) F1 score evaluations under linear-probing configuration. ViTAMIn-O demonstrates superior discriminatory capabilities in eight out of nine evaluated classes compared to DINOv2, Swin, and ViT baselines. (C) Accuracy comparisons of the fine-tuned ViTAMIn-O model against previously published domain specific benchmarks. ViTAMIn-O establishes new state of the art performance on the intestinal organoid dataset (surpassing CLORG, ViT and Tellu) and the hypothalamic-pituitary dataset (surpassing the EfficientNetV2-S and ViT ensemble model). On the colon budding and opacity datasets, the ViTAMIn-O model achieved performance parity in comparison to the D-CryptO model. Data points are either presented individually (C) or as mean ± standard deviation (A and B) across five independent runs. Across all runs, we selected a batch size of 32 (batch_size = 32) and epoch size of 20 (epochs = 20). Statistical analyses were carried out using Welch’s t-test.

**Figure 3.**
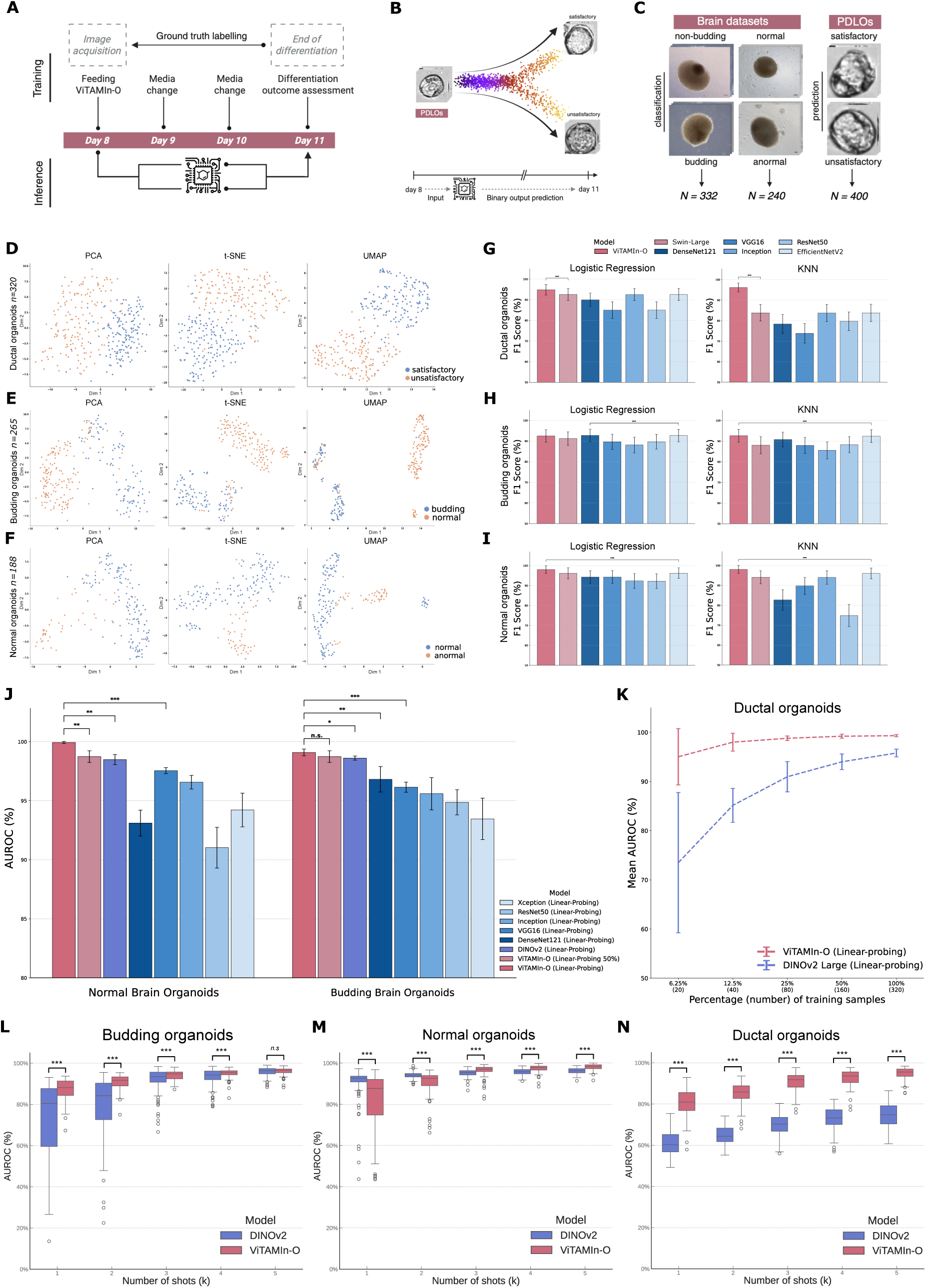
Unsupervised feature representation, external transferability, and data efficiency of ViTAMIn-O. (A) Schematic of the longitudinal prediction framework for PDLOs. Endpoint binary morphological outcomes (satisfactory and unsatisfactory), evaluated at day 11, were retroactively mapped to single-organoid images acquired at day 8. (B) Morphological divergence during PDLO differentiation. PDLOs transition from a homogenous baseline state at day 8 to heterogenous states by day 11. Vitamin-O leverages this to predict the binary day 1. Differentiation outcomes using solely day 8 imaging sets. (C) Representative transmitted light microscopy images of brain and PDLO models. Here, the experimental setup and downstream analysis distinguished between longitudinal prediction of future developmental trajectories of PDLOs and standard cross-sectional classification of morphological states in brain organoid models. (D-F) Dimensionality reduction visualizations including PCA (left), t-SNE (middle), and UMAP (right) of frozen ViTAMIn-O feature vector embeddings on never-before-seen datasets: PDLO, brain budding, and brain normal organoids. ViTAMIn-O natively clusters distinct morphological groups without pretraining. (G-I) Per-organoid class F1 scores demonstrating the linear separability of extracted features on brain and PDLO organoids. Models (ViTAMIn-O, Swin, DenseNet121, VGG16, Inception, ResNet50, EfficientNetV2) were trained exclusively using computationally lightweight machine learning algorithms, namely logistic regression and KNN. **j**, Linear-probing performance on the brain normal and budding datasets. ViTAMIn-O utilizing only 50% of the training data remains highly competitive against baseline models utilizing 100% of the data, with DINOv2 serving as the second best performing. (K) Performance titration on the external PDLO organoid dataset. ViTAMIn-O consistently outperforms DINOv2 in data sparse scenarios, maintaining robust predictive accuracy even when training data is artificially linearly reduced from 100% to 6.25%. (L-N) Few-shot transfer learning scenarios comparing ViTAMIn-O against DINOv2 across 1-shot to 5-shot settings. Data are presented as the mean across 100 experimental replicas, highlighting ViTAMIn-O’s superior adaptation capabilities under limited data regimes.

**Figure 4.**
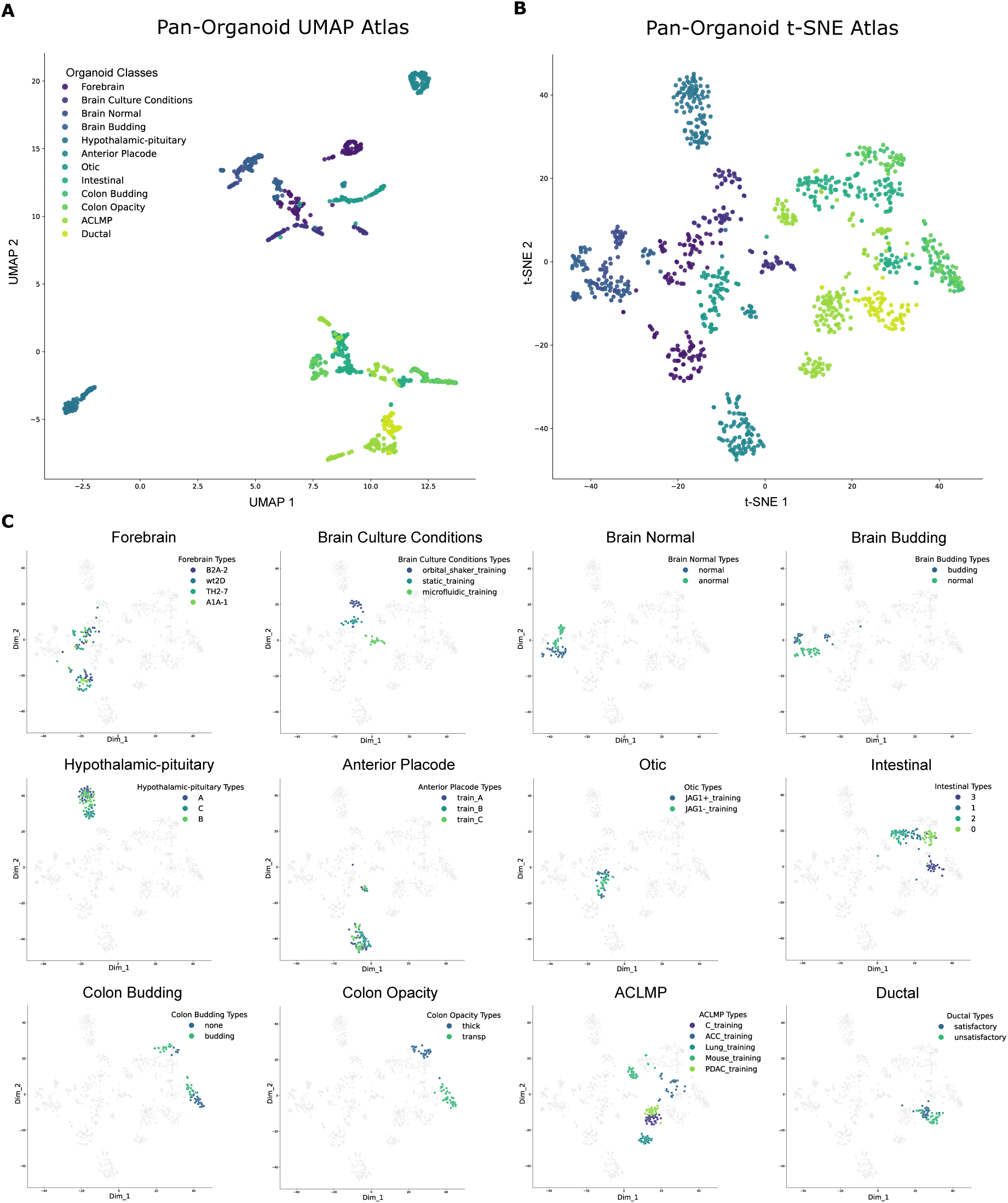
A comprehensive morphological atlas across organoid models reveals global lineage separation and local biological trajectories. (A) Global UMAP projection of ViTAMIn-O’s feature embeddings encompassing all 12 evaluated organoid datasets. The map naturally segregates along developmental germ layers, with endodermal associated tissues (e.g., intestinal or colon sets) on the bottom and ectodermal associated tissues (e.g., brain datasets) on the top. (B) Global t-SNE projection of the identical feature space, emphasizing the localized clustering and internal heterogeneity of the individual organoid classes. The number of individual points is normalized across all categories to balance the quantitative heterogeneity. (C) 12 fine grained, per-class t-SNE projections highlighting specific developmental and molecular trajectories against the grey background of the global dataset. Associated hyperparameters needed to derive the presented maps were generated using 20 nearest neighbor (n_neighbors = 20), a minimum distance of 0.1 (min_dist = 0.1), and the cosine distance metric for UMAP. For t-SNE, a perplexity of 30 (perplexity = 30) as well as 1,000 iterations (iterations = 1,000) were utilized.

**Figure 5.**
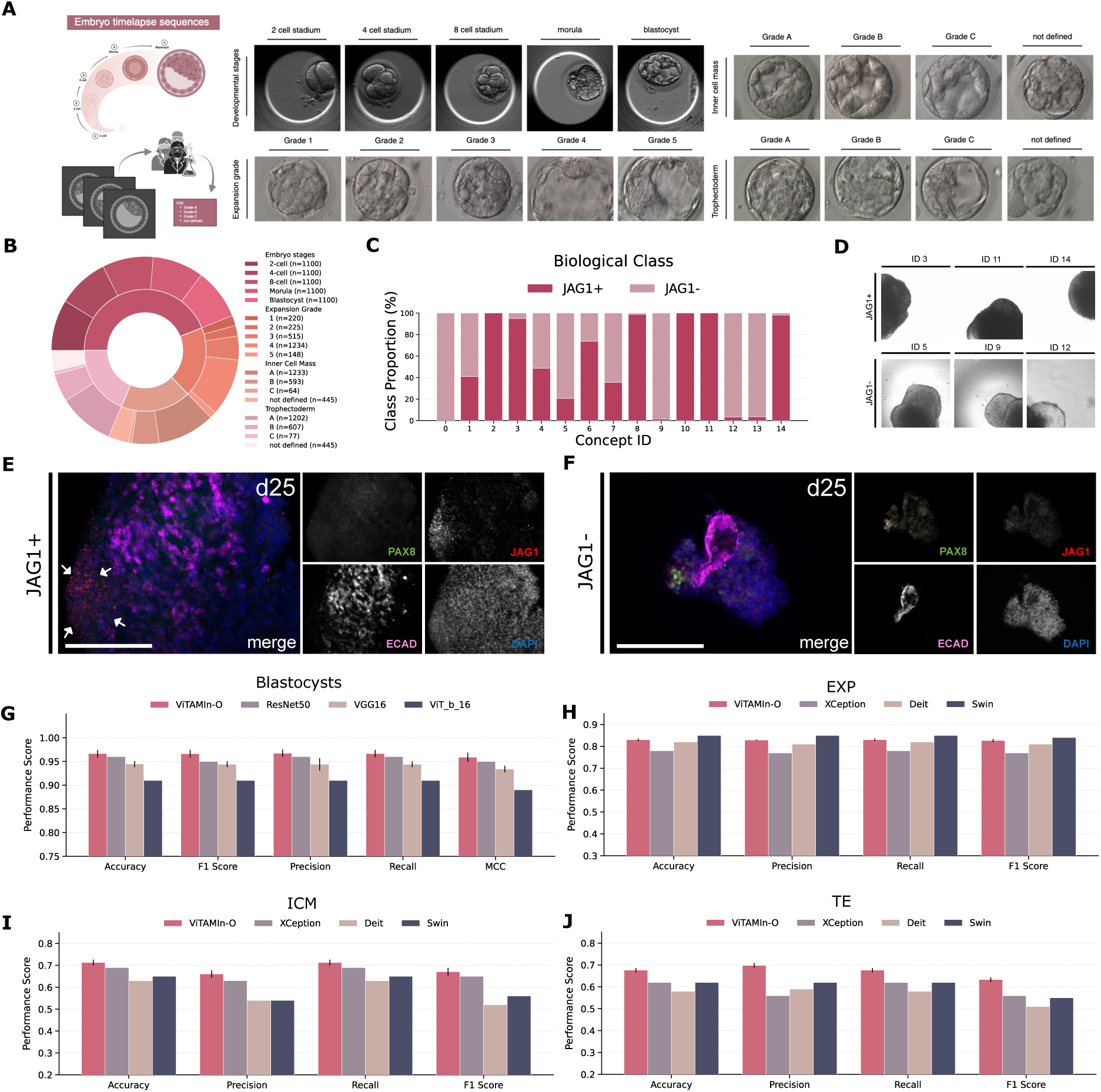
ViTAMIn-O integrates automated concept extraction (OrgACE) and demonstrates generalist adaptability to embryonic developmental staging. (A) Schematic overview of the expert-annotated embryonic transmitted light microscopy datasets. Embryos are categorized either by their concurrent developmental stage (2-cell to blastocyst) or by clinical Gardner grading criteria, assessing expansion grade, inner cell mass, and trophectoderm quality. (B) Composition of the test datasets. The inner ring visualizes the relative contribution of overarching global datasets (e.g., embryo stages, expansion grade), while the outer ring details the specific per-class distributions (e.g., 2-cell, 4-cell). Absolute quantitative shares for each category are provided on the right. (C-F), Automated Concept-based Explanation on Organoids (OrgACE) pipeline. (C) Normalized distribution of morphological concepts associated with JAG1 expression in otic organoids. (D) Representative brightfield images of algorithmically extracted concepts. (E) and (F), Immunofluorescence stainings demonstrating the presence (E) or absence (F) of JAG1 protein expression corresponding to the identified morphological signatures. (G) Benchmarking of the fine-tuned ViTAMIn-O model against established architectures (ResNet50, VGG16, ViT-B/16) on the classification of early expert assessed embryo stages^49^. (H-J) Comparative evaluation of ViTAMIn-O against XCeption, Deit, and Swin baselines on blastocyst quality classification^48^, encompassing (H), expansion grade (EXP), (I) inner cell mass (ICM), and (J), trophectoderm (TE) quality. Data are presented as mean ± standard deviation, obtained across five independent runs for the ViTAMIn-O architecture.

### Overview

The subsequent sections illustrate ViTAMIn-O’s capabilities in the context of fine-tuning, linear-probing and few-shot transfer learning, across a range of classification and prediction tasks. The process of fine-tuning refers to leveraging a model’s maximal separability capabilities, unfreezing of the model’s backbone encoder and the specification of a new classification layer for the designated task. In contrast, the linear-probing configuration trains a new classification layer on top of the frozen feature vector embeddings of the backbone encoder and relies on the extracted knowledge from pretraining. Linear regression classification (LRC) and k-nearest neighbour (KNN), simple forms of machine learning algorithms, rely on the feature vector embeddings of each image for a given dataset and subsequently discriminate the individual classes into the emerging categories, thereby even underscoring the computational costs of linear-probing. Few-shot transfer learning, in addition to that, utilises only a small percentage of the available training data under either fine-tuning, or more classically, linear-probing configurations.

To quantitively assess the performance and conduct extensive benchmarking comparisons between different models, we applied the following metrics in our analysis: Area under the curve receiver operating characteristic (AUROC), accuracy, precision, recall, F1 score (F1) and Matthew’s correlation coefficient (MCC). Means of statistical significance were assessed using the P values from the two-sided independent t-test with 1,000 resampled test sets if not noted otherwise.

### Comparative analysis on the hold-out test set

Presenting a designated platform for lab-internal deployments of highly specialized computer vision models requires robust transferability of the generalist model across a wide range of prediction tasks. To rigorously benchmark ViTAMIn-O against various organoid morphologies, we performed an internal evaluation on the hold-out test dataset secluded from the pretraining set. This evaluation encompassed 28 individual tasks summarized into nine global categories.

We first evaluated the feature extraction capabilities of ViTAMIn-O without updating the backbone network by employing a linear-probing configuration. We compared its performance against three state-of-the-art vision models, namely: DINOv2, Swin, and ViT (Figures 2A and 2B). ViTAMIn-O demonstrated striking robustness, outperforming the next best baseline models in all nine out of nine tasks evaluated by the Area Under the Receiver Operating Characteristic (AUROC) value (Figure 2A). Notably, ViTAMIn-O achieved substantial performance gains on tasks such as predicting the JAG1 expression in otic organoids, here summarized as otic (0.875 ± 0.022 vs. Swin: 0.628 ± 0.065), class separation on forebrain organoids (0.956 ± 0.002 vs. DINOv2: 0.857 ± 0.009), and intestinal organoids (0.983 ± 0.000 vs. DINOv2: 0.963 ± 0.001). This superiority was further reflected in the F1 scores, where ViTAMIn-O exceeded baseline models in eight out of nine tasks, with particularly clear margins in the anterior placode (0.613 ± 0.024 vs. DINOv2: 0.483 ± 0.053) or hypothalamic-pituitary organoid categories (0.724 ± 0.012 vs. ViT: 0.658 ± 0.010). DINOv2 only marginally outperformed ViTAMIn-O in the colon budding F1 score (0.899 ± 0.007 vs. 0.890 ± 0.005) (Figure 2B).

Protein expression patterns are key drivers of the functional and morphological state of cells as well as their cellular neighbourhood. If follows that a generalist model should be able to map the morphological features extracted from imaging data to the underlying biological states of the organoid. To this end, we conducted a finer grained per-class analysis on the hypothalamic-pituitary organoid dataset, predicting RAX expression levels: category A (70 < %RAX), category B (40 ≤ %RAX < 70), and category C (%RAX < 40). Strikingly, even under linear-probing, ViTAMIn-O achieved AUROCs of 0.903 ± 0.003, 0.786 ± 0.008, and 0.949 ± 0.002 across categories A, B, and C, respectively. These linear-probing results surpassed a previously published, highly specialized fine-tuned ensemble model, which achieved 0.878, 0.745, and 0.941 on the matched categories. Full fine-tuning of ViTAMIn-O further elevated performance in the corresponding intermediate and low expression classes (category B: 0.802 ± 0.011; category C: 0.952 ± 0.006), whilst preserving a strong baseline in the high expression class prediction (category A: 0.899 ± 0.007).

Finally, to situate ViTAMIn-O within the current landscape of specialized vision-based models, we contrasted the performance of our generalist model, applicable across diverse datasets, against previously published models restricted for only a single organoid class (Figure 2C). ViTAMIn-O established a new state of the art on the intestinal dataset, yielding an accuracy of 91.5% ± 0.4%, cleanly surpassing recent benchmarks including CLORG^56^ (86.9%), ViT^57^ (87.0%), and Tellu^43^ (83.6%). Similarly, on the overall hypothalamic-pituitary dataset, ViTAMIn-O significantly outperformed the Asano ensemble model of EfficientNetV2-S and ViT (accuracies of 74.6% ± 1.4% vs. 70.0%). On the D-CryptO tasks, the ViTAMIn-O model achieved performance parity with the D-CryptO architecture, recording a balanced accuracy of 90.6% ± 0.9% (vs. 90.9%) on colon budding and 98.0% ± 0.4% (vs. 98.0%) on colon opacity separation tasks. Taken together, these data illustrate that diversifying the pretraining data pool enables a single generalist model to not only incentivize morphological intricacies across various tissues but to match or surpass single task models engineered for biological imaging on organoids.

### External transferability and robust feature representation in data sparse environments

Real world deployment of deep learning frameworks in wet-lab environments necessitates robust transferability to entirely novel datasets not encountered during pretraining. Current computer vision models applied to organoid imaging are frequently restricted to singular tissue types or specific imaging modalities^58,59^, requiring extensive retraining whenever experimental protocols are altered. A true generalist model should possess feature vector embeddings structurally rich enough to capture universal morphological cues, allowing it to rapidly generalize to unseen scenarios even when relying on computationally lightweight algorithms or strictly limited data.

To rigorously evaluate the feature extraction capabilities of ViTAMIn-O, we probed its feature representation space using three external, previously unseen datasets: the normal brain, budding brain organoid sets, and one of the three newly generated imaging datasets used for this study, namely the pancreatic duct-like organoid (PDLO or ductal) imaging set. In brief, ViTAMIn-O was evaluated on its ability to predict developmental outcomes of PDLOs. By training the model on day 8 images retroactively paired with their definitive day 11 outcome labels, we probed its capacity to predict PDLO viability three days prior to the differentiation endpoint (Figures 3A and 3B). In contrast to the longitudinal forecasting for PDLOs, ViTAMIn-O was further evaluated on cross-sectional image classification for the brain organoid datasets. Here, by utilizing expert-annotated ground truth labels, the model was tasked categorizing current morphological states, distinguishing between normal and aberrant phenotypes. (Figure 3C)

Generating an intuitive understanding of a model’s high dimensional representations is essential for validating its zero-shot generalization. To this end, we visualized raw feature vector embeddings without any downstream training or fine-tuning using Principal Component Analysis (PCA), t-Distributed Stochastic Neighbor Embedding (t-SNE), and Uniform Manifold Approximation and Projection (UMAP) (Figures 3D–3F). Across all three dimensionality reduction techniques, ViTAMIn-O natively organized the unseen imaging data into highly separable, biologically coherent clusters. This inherent spatial segregation illustrates how the model successfully abstracted high dimensional, nonlinear morphological relationships during pretraining, negating the need for dataset specific retraining to achieve basic structural understanding of the rich semantic space of biologically informed morphological variability.

Because high quality feature embeddings should be linearly separable, we hypothesized that computationally inexpensive machine learning algorithms would suffice to achieve high predictive accuracy on these external datasets. We trained Logistic Regression Classifiers (LRC) and k-Nearest Neighbor (KNN) algorithms on the frozen ViTAMIn-O embeddings and benchmarked the per-class F1 scores against a variety of architectures (Figures 3G–3I). When leveraging a KNN algorithm, ViTAMIn-O consistently outperformed all baselines, achieving F1 scores of 98.05 ± 1.98% on the normal brain dataset (vs. EfficientNetV2: 96.00 ± 2.73%), 95.57 ± 2.53% on the budding dataset (vs. EfficientNetV2: 92.45 ± 3.10%), and 93.66 ± 2.83% on the PDLO dataset (vs. Swin-Large: 83.84 ± 4.04%). Under LRC configurations, ViTAMIn-O secured the highest performance on the PDLO dataset (94.92 ± 2.50% vs. EfficientNetV2: 92.53 ± 2.90%) and maintained performance parity on the normal and budding brain datasets, narrowly trailing convolutional neural networks like EfficientNetV2 (96.05 ± 2.76% vs. 96.22 ± 2.65%) and DenseNet121 (92.49 ± 3.03% vs. 92.75 ± 3.01%), respectively. These further underscores how ViTAMIn-O’s generalized representations allow basic algorithms to achieve clear discrimination on unseen organoid morphologies. To further mimic the reality of many biological laboratories, where the accumulation of quantitatively massive, annotated datasets remains costly and challenging, we evaluated ViTAMIn-O in a range of data sparse scenarios (Figures 3J–3K). Using linear-probing, ViTAMIn-O trained on only 50% of the available data maintained high AUROC scores on the normal brain (0.996 ± 0.001) and budding datasets (0.987 ± 0.005), closely matching its own performance being trained on the full 100% of the datasets (0.999 ± 0.001 and 0.991 ± 0.003, respectively) and surpassing the fully trained baselines (Figure 3J). We then moved forward and titrated the PDLO organoid training data down to progressively smaller fractions and compared ViTAMIn-O directly against DINOv2, the strongest competing baseline of the brain organoid classification comparison. Strikingly, when exposed to 6.25% (with n = 10 single organoid images per class, totalling n = 20 images) of the training data, ViTAMIn-O exhibited predictive superiority and narrower standard deviations compared to DINOv2 (0.950 ± 0.057 vs. 0.735 ± 0.143). This performance gap was maintained across the 12.5% (a total of n = 40 organoids, 0.980 ± 0.018 vs. 0.852 ± 0.035), 25% (n = 80 organoids, 0.988 ± 0.005 vs. 0.910 ± 0.031), 50% (n = 160 organoids, 0.992 ± 0.004 vs. 0.940 ± 0.016), and 100% (n = 320 organoids, 0.993 ± 0.002 vs. 0.958 ± 0.008) thresholds (Figure 3K).

Finally, to quantify the handling abilities of model limitations on data efficiency, we deployed ViTAMIn-O and DINOv2 in few-shot learning scenarios whereby models are exposed to as few as one training image per category, referred to as shots in the following section (Figures 3L–3N). Across 15 distinct few-shot scenarios (1-shot to 5-shot), ViTAMIn-O outperformed DINOv2 a total of 13 times, with 12 of these being statistically significant. The adaptability of ViTAMIn-O was particularly evident in the prediction of the differentiation outcomes of PDLO organoids, where it vastly outperformed DINOv2 at both the 1-shot (0.804 ± 0.066 vs. 0.608 ± 0.056) and 5-shot marks (0.948 ± 0.027 vs. 0.747 ± 0.062). Similarly, on the brain budding dataset, ViTAMIn-O led decisively in 1-shot performance (0.871 ± 0.052 vs. 0.725 ± 0.183). While DINOv2 displayed a statistically significant advantage in the 1- and 2-shot settings of the normal brain dataset (1-shot: 0.903 ± 0.086 vs. 0.813 ± 0.142), ViTAMIn-O’s performance recovered by the 3-shot mark up to the 5-shot mark (0.981 ± 0.015 vs. 0.962 ± 0.014). Collectively, these results reinforce how the diversification of the pretraining data pool enables ViTAMIn-O to transfer its separability capabilities under novel experimental conditions, effectively underscoring its utility in real world deployment even in data scarce environments.

### Unsupervised dimensionality reduction techniques illustrate how ViTAMIn-O internalizes developmental lineages through morphological atlases

In-depth profiling of organoids frequently requires functional readout, imaging modalities and omics technologies. While these techniques have provided precise molecular atlases of organoid development, they remain costly and mostly static. A precise definition of optimal timepoints to halt differentiation and perform these analyses would be therefore valuable. A purely vision-based morphological atlas cannot only provide a cost-efficient alternative to extract underlying biological signals but also to define optimal timepoints enabling researchers to extract underlying biological signals, identify optimal temporal windows to halt differentiation and ultimately track continuous cellular trajectories through integration of single-cell omics techniques. To determine whether ViTAMIn-O is able to construct a topological atlas, we performed dimensionality reduction and projected the feature embeddings across all nine pretraining and three external datasets into a unified morphological atlas (Figure 4).

We first utilized Uniform Manifold Approximation and Projection (UMAP) to visualize the entire dataset of this study, which aims to preserve global topological relationships in a nonlinear way (Figure 4A). Strikingly, the UMAP projection generally segregated the organoid model systems into two broad, biologically distinct hemispheres. Tissues associated with the endodermal lineage, including intestinal, colon budding, colon opacity, ACLMP and additional external PDLO dataset, were anchored on one side of the embedding space. Conversely, tissue cultures associated with the ectodermal or more specifically neural lineage, encompassing forebrain, normal brain, budding brain, hypothalamic-pituitary, anterior placode and otic organoids were positioned on the opposing upper hemisphere. This topological separation indicates that ViTAMIn-O may have internalized the knowledge to distinguish the foundational morphometric signatures of distinct embryonic lineages in an unsupervised way.

Complementing this UMAP visualization, a global t-distributed Stochastic Neighbor Embedding (t-SNE) projection (Figure 4B) further clustered the global and local neighbourhoods, highlighting the dense internal heterogeneity within these respective organoid classes. To further extract biological insights from these local manifolds, we constructed 12 fine grained compartmentalized t-SNE maps (Figure 4C) based on the pan-organoid t-SNE projection (Figure 4B). These sub-projections illuminate distinct developmental dynamics or morphological states that are critical for quality control of the respective organoid cultures. For instance, the intestinal organoid manifold delineates a structural bifurcation, starting from baseline spheroids (category 0), the embeddings naturally map a divergence either towards a healthy developmental vector of early and late organoids (categories 1 and 2) or towards an aberrant cystic phenotype (category 3). By monitoring the positional information of an organoid batch within this manifold, researchers can non-invasively identify ideal differentiation endpoints or selectively exclude dysmorphic batches early on during differentiation (Supplementary Figure 4B).

Crucially, the per-class embeddings further demonstrate the model’s proficiency in linking morphological cues to the underlying molecular identity. The hypothalamic-pituitary embeddings did not form discrete categories but rather organize into a continuous spatial gradient that faithfully correlates with the continuous whole organoid RAX protein fluorescence intensity, whilst being annotated in a binary manner to facilitate the adoption of an appropriate classification algorithm. Conversely, in the brain organoid culture condition t-SNE plot, the spatial embeddings of organoid populations cultured in orbital shakers, static or microfluidic devices, were separated rather clearly, effectively matching organoid populations to environmental changes. Remarkably, this capacity for temporal and developmental inference extends to zero-shot scenarios. When applied to a holdout dataset of human preimplantation embryogenesis, the model naturally reconstructed the time adherent developmental sequence from the 2-cell to the blastocyst stage without any prior pretraining (Supplementary Figure 4C). Ultimately, these visualizations demonstrate that ViTAMIn-O offers a robust, non-destructive coordinate system for modelling stem cell state and fate, going beyond mere image classification.

### Automated concept-based explanations link macroscopic morphology to molecular states

While deep learning architectures can achieve high predictive accuracy in biomedical imaging, their black box nature limits their utility for mechanistic biological explanations to derive novel or confirm preexisting hypotheses. Conventional explainable AI methods, such as pixel attribution heatmaps (e.g., Grad-CAM), highlight localized regions within single images but fail to provide global, biologically actionable features that wet-lab experimentalists can consistently track across experimental batches in a high throughput way. To bridge the gap between computational embeddings and interpretable biology, we employed OrgACE to extract and quantify macroscopic, invariant morphological concepts.

To evaluate the utility of this framework, we applied ViTAMIn-O and the OrgACE algorithm to decipher the morphological correlates of JAG1 protein expression as a key developmental marker, indicative of the successful differentiation in the otic organoid model system. ViTAMIn-O was fine-tuned on the binary dataset, whilst the imaging data was subsequently divided into predefined patches, measuring equal proportions of the original images and mapped through a k-means clustering algorithm operating within the latent space of the fine-tuned ViTAMIn-O model. This unsupervised process automatically clustered the morphological patches into 15 distinct, interpretable concepts, and quantified their proportional association with the underlying JAG1 expression states, upon which the model drives its predictions (Figure 5C).

These series of steps identified highly specific phenotypical signatures linked to the underlying molecular identity associated with JAG1 protein expression. For instance, concept 3 demonstrated a 95% association with the JAG1+ state, whereas the appearance of concept 5 was associated with the absence of JAG1-expression. By extracting representative images for these algorithmic clusters, users can visually define the specific structural motifs driving the model’s predictions (Figure 5D). Concepts associated with JAG1-organoids (Concepts 5, 9, and 12) frequently featured distinct morphological signatures, such as the appearance of translucent cyst structures, localized at the outer rim of each organoid, absent in the primary JAG1+ clusters. These, in turn, represented themselves with tightly packed, denser edges (Concepts 5, 9, and 12).

To ground these computationally derived morphological concepts in a verifiable way, we performed immunofluorescence stainings, covering key developmental marker proteins in otic organoid differentiation^47^. Immunofluorescence analysis of day 25 otic organoids supported the structural concepts identified by OrgACE, where organoids displaying the more pronounced macroscopic features of the JAG1+ concepts exhibited localized JAG1 protein expression strictly at the outer organoid rim (Figure 5E; Supplementary Figure 3E). Conversely, organoids exhibiting the JAG1-morphological concepts lacked this spatial fluorescence signal entirely (Figure 5F; Supplementary Figure 3F).

Collectively, these results illustrate how the ColabViTAMIn-O platform enables researchers to translate AI-guided predictions into testable, morphologically guided hypotheses, establishing a closed loop framework between non-destructive brightfield imaging, computational prediction, and spatial molecular validation.

### Generalization to pre-implantation embryonic development and morphological grading

A key limitation of several current biomedical computer vision tools is their strict domain specialization. Algorithms trained on specific in vitro morphologies frequently fail when translated to related, yet distinct, model systems. If a model is to serve as a genuine generalist tool for stem cell biology, its underlying feature representations should translate reliably to developmental models other than organoids, such as human embryo imaging datasets. To evaluate ViTAMIn-O’s adaptability to external, non-organoid or more generally speaking, stem cell domains, we fine-tuned and benchmarked the model on two independently published, expert-annotated embryo imaging datasets encompassing early developmental staging and blastocyst quality grading.

We first assessed the model’s capacity to classify early developmental transitions using a dataset of 5,500 embryo images categorized into 2-cell, 4-cell, 8-cell, morula, and blastocyst stages^49^. ViTAMIn-O was fine-tuned and evaluated against the established baseline architectures for this dataset, including ResNet50, VGG16, and ViT-B/16 (Figure 5G). ViTAMIn-O demonstrated consistent predictive superiority and robustness across all five evaluation metrics, achieving an accuracy of 96.68 ± 0.77%, an F1-score of 96.65 ± 0.79%, and a Matthews Correlation Coefficient (MCC) of 95.88 ± 0.96%. These results marginally exceeded the second best performing recently published benchmark of ResNet50 (accuracy: 96.0%; F1-score: 95.0%; MCC: 95.0%), indicating that ViTAMIn-O’s feature embedding space is applicable to embryonic cleavage events.

To test the model on rather nuanced, clinically relevant morphological assessments, we applied ViTAMIn-O to a second dataset of 2,344 blastocyst images^48^. This dataset requires the classification of three distinct morphological quality metrics, namely the expansion grade (EXP), inner cell mass (ICM), and trophectoderm quality (TE). We benchmarked ViTAMIn-O against published benchmarks of XCeption, Deit, and Swin architectures (Figures 5H–5J). On the expansion grading task, ViTAMIn-O achieved highly competitive performance (accuracy: 83.02 ± 0.55%; F1-score: 82.87 ± 0.23%), maintaining close performance parity with the Swin architecture, which served as the leading model for this specific task (accuracy: 85.0%; F1-score: 85.0%).

Importantly, ViTAMIn-O demonstrated a distinct advantage in the tasks of ICM and TE grading. Across all models which were evaluated in this regard, these category tasks yielded the overall lowest performance metrics illustrating their inherent complexity. For ICM classification, ViTAMIn-O outperformed the next best baseline, XCeption, yielding an accuracy of 71.30 ± 1.29% (vs. 69.0%) and an F1-score of 66.07 ± 1.69% (vs. 63.0%). This performance advantage was further extended in TE classification, where ViTAMIn-O achieved an accuracy of 67.64 ± 0.97% and an F1-score of 69.81 ± 1.05%, surpassing the baseline Swin and XCeption models (accuracies: 62.0%; F1-scores: 62.0% and 56.0%, respectively). Collectively, these evaluations underscore how the pretraining regimen of ViTAMIn-O yields a highly versatile visual foundation capable of adapting to complex, stem cell models without requiring architectural redesigns.

## DISCUSSION

In this article, we introduce ViTAMIn-O, a generalist computer vision foundation model specifically designed to decode stem cell and organoid-based model systems as well as their morphological trajectories, alongside its dedicated deployment platform, ColabViTAMIn-O. Our approach departs from the standard paradigm of training narrow, task specific classifiers. Instead, we extracted and harmonized feature representations across diverse organoid datasets, spanning multiple tissue lineages and heterogeneous imaging modalities, into a single transformer-based architecture. Through extensive benchmarking evaluations, encompassing zero-shot feature extraction, data scarcity simulations, and automated concept generation, we demonstrate how a model pretrained on a vastly diverse morphological dataset consistently achieves superior generalization compared to models tailored to singular tasks. These results illustrate that biological data heterogeneity can serve as a key accelerator, rather than a barrier, for developing the next generation of robust, widely deployable computational tools for tissue engineering and developmental biology.

ViTAMIn-O establishes a new state of the art in the assessment of organoid differentiation by aggregating diverse morphological information during pretraining. The model’s robust performance, consistently outperforming leading vision foundation models such as DINOv2^60^, Swin^50^, and ViT^13^ across rigorous linear-probing evaluations, underscores the critical importance of leveraging diverse, lab-internal datasets to harness the full discriminative power of deep learning. ViTAMIn-O’s domain expertise translates directly to highly specialized tasks. It achieved pronounced, multi metric increases over previously published benchmarks on the intestinal dataset (surpassing CLORG^56^, Tellu^43^, and ViT^57^) and the hypothalamic-pituitary dataset (surpassing a specialized ensemble architecture^35^). Furthermore, it closely matched the performance of the D-CryptO^44^ model on colon datasets, emphasizing how a single CV model can effectively replace multiple isolated, single use algorithms in the domain of various stem cell model systems. Crucially, ViTAMIn-O exhibits robust domain adaptation to never before seen datasets, maintaining robust predictive performance even in environments simulating extreme data scarcity. By isolating the model’s feature vector embeddings, we demonstrated that computationally lightweight algorithms, such as k-nearest neighbor, are sufficient to achieve state of the art discrimination on external PDLO and brain organoid datasets. By iteratively halving the training set for the prediction of PDLO organoid differentiation, ViTAMIn-O consistently maintained its predictive advantage, exhibiting highly robust AUROC scores even when exposed to as little as 6.25% of the available data. This data efficient utility was further shown in few-shot learning scenarios, where ViTAMIn-O secured performance advantages over DINOv2 using only 1 to 5 image samples per category across all scenarios. These insights further summarize how the internal representations of ViTAMIn-O are of exceptional quality, supporting rapid knowledge transfer across diverse morphological domains.

This rich feature space extends beyond classification tasks, enabling the unsupervised reconstruction of morphologically grounded feature maps. By projecting the high dimensional embeddings into a unified morphological atlas, we demonstrate that ViTAMIn-O natively segregates distinct embryonic germ layers, autonomously separating endoderm associated from ectoderm and neural associated tissue cultures. Moreover, these latent embeddings capture continuous spatio-temporal dynamics, accurately mapping the morphological bifurcations of intestinal organoids and the continuous protein expression gradients of hypothalamic-pituitary cultures. Remarkably, this capacity for temporal inference was found to be generalizable to human embryonic development stages. ViTAMIn-O successfully reconstructed the chronological cleavage stages of mammalian embryos and achieved highly competitive performance in blastocyst quality grading, underscoring its versatility as a universal tool for stem cell biology.

To ensure these advanced computational capabilities translate into tangible wet-lab utility, we integrated these functions into ColabViTAMIn-O. A primary bottleneck in the adoption of versatile AI tools in biology is the requirement for advanced programming expertise and specialized hardware. ColabViTAMIn-O addresses this by providing an intuitive, completely code free platform hosted on a cloud infrastructure. Through this interface, researchers with no prior machine learning experience can execute tasks such as linear-probing, few-shot transfer learning, fine-tuning and predictive model evaluation, using their own lab-internal datasets. Furthermore, the platform integrates the Automated Concept-based Explanation on Organoids (OrgACE) pipeline. By automatically clustering morphological patches and linking them to molecular states (e.g., JAG1 expression in otic organoids), OrgACE allows experimentalists to derive undisruptive, morphologically guided hypotheses directly from the ColabViTAMIn-O interface, establishing a closed loop cycle of imaging, prediction, and henceforth, discovery. Finally, we introduce the ViTAMIn-O Model Hub, a centralized repository integrated within the ColabViTAMIn-O infrastructure. This platform allows researchers to upload, access, and deploy trained models, facilitating both individual research applications and broader community-wide collaboration.

Despite the highly robust performance that ViTAMIn-O demonstrates, several conceptual and technical challenges remain. Primarily, vision-based models, such as the current ViTAMIn-O architecture, rely entirely on macroscopic morphological signals, derived from variable imaging settings, which serve only as proxy indicators for underlying gene and protein regulatory networks. While OrgACE successfully links structural concepts to targeted protein expression, pure computer vision cannot mechanistically explain the complex transcriptomic drivers of cell fate trajectories. To derive deeper insights into developmental processes, future iterations of this framework must evolve into multimodal architectures. By correlating non-invasive visual modalities with spatially resolved molecular data and single-cell omics sequencing efforts, we aim to directly map the latent morphological space onto proteomic as well as transcriptomic landscapes. Additionally, while the accumulation of diverse imaging datasets establishes ViTAMIn-O as a generalist foundation model, real world deployments across an even broader range of uncharacterized organoid classes and dynamic time lapse imaging modalities will be essential to continually refine its generalization capabilities for future use cases.

In conclusion, we present ViTAMIn-O and ColabViTAMIn-O as a unified, generalist AI ecosystem designed for stem cell and organoid biology. The model’s robust performance is driven by the premise, that diverse developmental models share common, domain invariant morphological features that can be computationally extracted and transferred across distinct tissue classes. Considering the vast, untapped repository of lab-internal bright-field and phase contrast imaging data globally available, ColabViTAMIn-O and ViTAMIn-O Model Hub provide an open source, code free framework for its immediate utilization and democratization. By releasing both the model weights, the aggregated training datasets, and the open platform, we aim to catalyse the democratization of AI in stem cell biology, accelerating protocol optimization and unlocking new avenues for non-invasive screening in future regenerative medicine efforts.

## RESOURCE AVAILABILITY

### Data and code availability

- All datasets generated for and used in this study will be made publicly available upon acceptance under the following link: https://zenodo.org/records/19466176?token=eyJhbGciOiJIUzUxMiJ9.eyJpZCI6ImRmMmM3YTdi LTVkZTYtNGZmMC04ZTk1LTA5NDI4NTY5MGZiZCIsImRhdGEiOnt9LCJyYW5kb20iOiJlYzg0YT JkMjJkYWIwMzdjNjFkNDU4YTU5OTY1MDZhMyJ9.Q1xrTzJ_YTHDOKEQKuATkQ6TiB6WjnQ-iwh8ofi2j-1X9sG5Wu_YElizhftTp6k88SfL3Qva3PAW5PpT_zfB0g.
- The ColabViTAMIn-O platform is accessible under: https://colab.research.google.com/drive/1nDXBvSziGndkVcpUH9jxRbZpwX-2MMDU?usp=sharing. The ViTAMIn-O Model Hub can be accessed through https://huggingface.co/ViTAMIn-O. All source code used for this study has been archived on Zenodo and is temporarily restricted for the process of peer review. Reviewers can access the code via the following link: https://zenodo.org/records/18942762?token=eyJhbGciOiJIUzUxMiJ9.eyJpZCI6IjQ4MGE3YzYwLTY5MDUtNDM4NC05OTA5LTU1ZGE1NDQ0ZWJjZiIsImRhdGEiOnt9LCJyYW5kb20iOiI5ODk0MjM3NjFjMzJjZjllOTBjNDU1YWE1M2NiNmUxZiJ9.-Wnd02sZyI1J4ZSC3pMcetLeCFrdZiCiNCsW4aQdjQBVkBc3Bwv-_aO_es46f4v1oG_Aa_iQd0SL2Wapk8NSnw. The pretrained ViTAMIn-O model is an open source model with a MIT license and is available on Zenodo via following link: https://zenodo.org/records/18771069?token=eyJhbGciOiJIUzUxMiJ9.eyJpZCI6ImExYTBhMzk5LTlmYTEtNDU2OC1iNGIwLWY5YWY5NjQxZDhjZiIsImRhdGEiOnt9LCJyYW5kb20iOiI2ZDRjNjVlNjlmM2JlMzBkNzk4MTIzMjVlMTkyZGM1MyJ9.t7Xg4gwgLthFVY7Yu_cG9Cs2-C3RdpHpxfcxk7K4H0u2XMBAmfYNUD8v-ANYVKZAsHBx_e1EtynQsDH_oD719Q or through the ViTAMIn-O Model Hub.
- Any additional information required to reanalyze the data reported in this paper is available from the lead contact upon request.

## ACKNOWLEDGMENTS

We would like to express our gratitude towards Sabine Conrad for her technical assistance. This work has been supported by the Friedrich Ebert Foundation as well as the IZKF Promotionskolleg (2024-2), Faculty of Medicine Tübingen, University of Tübingen to F.H. and by the DFG: LI 2044/7-1, LI 2044/5-1, LI 2044/5-2 to S.L.. We acknowledge the support from the Open Access Publishing Fund of the University of Tübingen.

## AUTHOR CONTRIBUTIONS

F.H., S.K. and M.K. designed and contributed to the conception of the study. F.H. wrote the manuscript, created the schemata and figures, performed the experiments with the anterior placode organoids, acquired and organized the datasets, conducted the computational experiments, analysed and interpreted the data. B.B. performed experiments with otic organoids. A.T.K., J.L., and M.B. designed and A.T.K. performed the experiments with PDLO organoids. M.B., A.V., K.A., N.P., A.K., S.L., S.K. and M.K. critically revised the manuscript and provided guidance and key insights for the research. All authors contributed to the article and approved the submitted version.

## SUPPLEMENTAL INFORMATION

**Document S1. Figures S1–S5**

## METHOD DETAILS

### Training framework for ViTAMIn-O and knowledge distillation mechanism

ViTAMIn-O employs a teacher-student network-based architecture, leveraging knowledge distillation techniques specifically tailored for multi-task learning on heterogenous organoid imaging datasets. Under this premise, ViTAMIn-O draws inspiration from previously published methods in the domain of CV in large scale visual representation learning. At its foundation lays the objective of ensuring an efficient crosstalk between supervisory stability of the teacher and promoting feature robustness in the student.

The backbone encoder of both the teacher and student model was adopted from the Swin-Large Vision Transformer (“microsoft/swin-large-patch4-window7-224”)^50^. The decision to select the Swin architecture as the core of ViTAMIn-O was taken given its ability to process imaging data in a hierarchical manner using the shifted window attention which enables it to efficiently capture both the fine-grained morphological features in addition to rather global structural contexts of individual organoids. Crucially, both the teacher and student backbones were initialized with the published weights derived from training on the ImageNet-1k dataset at an input resolution of 224 x 224 pixels (https://huggingface.co/microsoft/swin-large-patch4-window7-224).

To incentivize a highly generalized feature space concerning organoid morphology while simultaneously preserving the capacity for task specific classification and prediction, we trained ViTAMIn-O with nine organoid Multi-Layer Perceptron (MLP) classification heads. These heads, corresponding to nine organoid imaging datasets, comprise a total of 28 individual classification classes. Global Average Pooling (GAP) was implemented to bridge the feature map output of our Swin-based models to the subsequent classification heads. To enable feature consistency distillation between the teacher and student model, dedicated projection heads were implemented. Precisely, training was facilitated by a pooled dataset of several brain organoid models, encompassing forebrain^41^, various brain culture conditions^42^, hypothalamic-pituitary^35^ as well as anterior placode^46^ and otic^47^, intestinal^43^, two colon^44^ and a pooled dataset of lung epithelia, human pancreatic ductal adenocarcinoma (PDAC), salivary adenoid cystic carcinoma (ACC), mouse small intestine and colon epithelia organoids (ACLMP for short)^45^. To ensure compatibility with the pre-trained backbone, the input data, including TIF, PNG and JPG formats, were pre-processed through a Swin-compatible auto image processor to an input resolution of 224 x 224 pixels. A dual data splitting protocol was implemented to handle the heterogeneity in quantitative dataset imbalances. In accordance with previously published benchmarking studies, the data points associated with the intestinal and colon opacity datasets were split into a predefined static separation into training, test and validation sets^43,57^. The remaining training sets were dynamically split using a 20% stratified validation split, ensuring the maintenance of the original class distribution between the generated training and validation sets whilst preventing sampling bias.

The core training data pipeline was implemented through a bifurcating system, where the input data was split, ensuring that the teacher maintains its reliable supervision for the student, which in turn, receives the necessary regularization for robustness in training. Precisely, one path fed the student model aggressively augmented images while the other stable, non-augmented path fed into the teacher. Specifically, the student path was designed in a way to promote the learning of highly relevant morphological features of organoids which are robust and invariant to photometric distortions. This extensive stochastic augmentation proved to be essential for the generalization of common features in heterogenous organoid classes and imaging modalities (Supplementary Fig. 5). Geometric transformations include quasi random resizing and cropping, flipping along the vertical and horizontal axis, rotating up to 30°. The pipeline further incorporates colour jittering and random erasing, which further promote the model’s invariance to structural and photometric inconsistencies across datasets.

The student model was trained cyclically across all nine datasets by implementing a specialized scheduling mechanism. In essence, the training pipeline utilized a batch level, task balanced round robin sampling strategy to ensure that within every step of training cycles, the model processes a small batch from each of the tasks sequentially. This cycling method prevents the model from employing disproportionate representation capacities to the largest or easiest task which would otherwise defy the purpose of aggregating this diverse imaging dataset. Instead, through this forced balancing in dedicating its computational capacities to all tasks, the model is allowed to learn a rather robust feature space and thus generalizes more effectively across all organoid types. The training utilized a maximum of 50 epochs, constrained by an early stopping mechanism with a patience of 10 epochs. Training termination was governed by the average student validation loss across all tasks, thus ensuring that the final model exhibits the overall best performing generalization capabilities rather than focusing on a single, potentially dominant class. Furthermore, the training was optimized through the Adam optimizer with differential learning rates applied to the backbone and the task specific heads to stabilize the process of feature extraction with a value of 1×10^-5^ for the backbone and 1×10^-4^ for the task-specific heads. A batch size of 64 was deemed most suitable for stable training the baseline ViTAMIn-O model on a single A100 GPU with 40 GB VRAM for approximately 24 hours.

The total loss function (L_total_), a multifactorial function balancing supervised learning and knowledge distillation from a stable teacher model (W_T_) for the student model (W_S_). The teacher’s weights (W_T_) are updated through Exponential Moving Average (EMA) of the student’s weights with a momentum of µ = 0.999, ensuring that the teacher supplements the student with smooth and gradually evolving supervisory signals.

The total loss function (l total) thus leverages a supervisory classification term (L_cls_) and a weighted consistency term (L_consist_):

Where the consistency term is defined as follows:

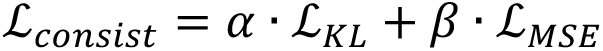

To address the class imbalances across the 28 sub classes, the Classification Loss (L_cls_) employs Class-Weighted Cross-Entropy. The consistency term (L_consist_) mixes the Kullback-Leibler (KL) Divergence (L_KL_) for logit matching with an alpha value (α) of 1.0 and the Mean Squared Error (L_MSE_) for feature level alignment with a beta value (*β*) of 0.25. To further maintain training stability, we linearly ramped up the consistency weight (w_consist_) over the first ten epochs.

### Datasets for training and evaluating ViTAMIn-O

We collected nine imaging datasets across six independent laboratories with a total of n = 32,773 images, divided into a training set of n = 24,751 individual organoid images as well as n = 3,611 and n = 4,411 for validation and internal testing respectively, n = 1,260 organoids for the external test set, additionally to a total of 7,844 annotated human pre-implantation embryo images. The models were tasked with predicting the differentiation endpoint outcomes across four datasets using two distinct approaches: either directly inferring longitudinal developmental trajectories for PDLO and anterior placode or_indirectly projecting underlying protein marker expression future differentiation successes in hypothalamic-pituitary and otic organoid datasets. The remaining datasets were utilized to evaluate binary or multiclass cross-sectional classification capabilities of models.

### Intestinal organoid dataset

The intestinal organoid dataset^43^ was generated by using a EVOS FL microscope (Thermo Fisher) with a 4x objective. A total of 840 individual whole plate images over the course of six days were acquired in total, resulting in an unequal distribution of organoid classes. Specifically, the authors assessed the organoids based off morphological criteria, corresponding to specific differentiation states. The training and subsequent testing datasets consist of category 0 (52%): early stage, small, cystic, non-budding, thick-walled organoids, category 1 (24%): early organoids with 1-2 crypts (buds), category 2 (14%): larger organoids with 3 or more crypts, and category 3 (10%): representing overstimulated, larger thin-walled organoids. Crucially, the classification task was to separate single organoids cross-sectional into one of the four aforementioned categories. It is important to note that crypt-derived intestinal organoids can either differentiate into budding organoid or spheroids, based off varying stochastic factors. All images were cropped and resized to 224 x 224 pixels^57^, as described by the authors. Datasets were pre-separated into a training set (n = 18,537), validation set (n = 2,058) and test set (n = 2,468). The datasets can be accessed through the following repositories: https://doi.org/10.5281/zenodo and https://doi.org/10.5281/zenodo.14725323).

### Hypothalamic-pituitary organoid dataset

Hypothalamic-pituitary organoids^44^ were imaged using an All-in-One Fluorescence microscope (KEYENCE BZ-X710). Images were acquired in the bright-field configuration using a 4x objective with a fixed exposure time of 1/3.5 s. Acquisition focused on the centre of the corresponding well with an invariant z-axis across all wells. The dataset consists of 1,500 images of individual organoids, with a training dataset of n = 1,200 and a test dataset n = 300, equally distributed into categories A, B and C. Organoids were grouped based on the relative organoid wide area expression of RAX protein on day 30 of differentiation: category A (70% < RAX), category B (40% ≤ RAX ≤ 70%) and category C (%RAX < 40) on day 30 of differentiation by experts. As such, models were tasked to predict the relative RAX expression levels derived only from bright-field imaging data of the subsequent organoids at day 30. Images were provided with input resolutions of 1,920 x 1,440 and were subsequently resized to 224 x 224 pixels. The data partitioning used in this study was adopted from the original authors and is available under https://doi.org/10.6084/m9.figshare.24616506.

### Forebrain organoid dataset

Forebrain organoids^41^ were generated through reprogramming peripheral blood mononuclear cells (PBMCs) of four patients into pluripotency. Four induced pluripotent stem cell (iPSC) lines of a healthy control (wt2D), two patients with TUBA1A- and TUBB2A-associated tubulinopathy (A1A and B2A) and one with tyrosine hydroxylase deficiency (TH) were generated. In total, 64 organoids were imaged in two independent laboratories over 12 timepoints (from day 2 until day 30), resulting in a total of 1,407 images (organoid 50, day 12, Lab A, after embedding was removed). Ultimately, models were tasked to classify organoid images across all timepoints into one of the four categories, solely according to their respective lineage (wt2D, A1A, B2A, and TH) illustrating their capabilities of lineage inference. Images were acquired with the Leica DMi 1 and Zeiss Axio Vert.A1 at 5x magnification. Images were split into a training set (n = 1,125 images) and test set (n = 282) with equally distributed data points between the four patient subfolders. Source data are publicly available under https://doi.org/10.5281/zenodo.10568828.

### Anterior placode organoid dataset

Olfactory placode organoids were imaged on day 12 and day 25 of differentiation, generating an equal batch of 100 images for three classes: undefined (category A), cystic (category B) and placodal (category C), featuring a total of n = 300 images. Single organoid level labels were assessed through expert-guided annotations on day 25 and retrospectively assigned to images acquired on day 12. Morphologically, on day 25, placodal organoids (satisfactory) display a translucent pseudostratified epithelium on their outer edges whilst preserving a denser core and a diameter of less than 300 µm. Cystic organoids (unsatisfactory), demonstrate larger diameters, homogenous edges, cores, outgrowths and an absence of an organized apical cell layer. To this end, the models were tasked to predict the endpoint differentiation outcome of single organoids at day 25 on the basis of imaging data acquired on day 12. A key caveat is that immunohistochemical studies must be performed to determine true cellular identity, as morphological insights only reveal correlated tendencies. Image acquisition was performed with the EVOS FL Colour Imaging System AMF 4300 in the phase contrast configuration (4/15 PH), 4x magnification at a resolution of 1,280 x 960 pixels with differing z-axis planes to generate high resolution images and account for well-to-well variability.

### Otic organoids

Otic organoids were generated from healthy human induced pluripotent stem cells (hiPSCs) by adapting previously established differentiation protocols^47,61,62^. In brief, hiPSCs were sequentially differentiated using E8, serum-free, as well as DFNB media, whilst being supplemented with specific growth factors and small molecules (including FGF 2/3/8/10/19, BMP4, CHIR-99021, EGF, IGF1, Y-27632, Wnt3a, heparin, BDNF, and NT3). For the imaging dataset, twelve individual organoids were tracked and imaged using the Evos FL Colour Imaging System AMF 4300 with a 4x magnification and resolution of 1,280 x 960 pixels. Brightfield images were acquired daily from day 10 up to day 25 of differentiation (16 continuous timepoints per organoid), yielding a total dataset of 192 images. Organoids were categorized into two distinct classes based on their subsequent immunofluorescence staining readout, JAG1 positive (n = 6) and JAG1-negative (n = 6). To rigorously assess model performance, the 192 images were divided into a training set (n = 154) and a holdout test set (n = 38), with the split being equally stratified to entirely omit class imbalances. To validate the morphologically derived computational concepts, endpoint immunofluorescence spatial stainings were performed. Organoids were fixed in 4% paraformaldehyde containing 10% sucrose, embedded in O.C.T. compound, and cryosectioned at 14 μm. Sections were then blocked and stained for the markers JAG1, PAX8, E-Cadherin (E-Cad), and DAPI. Optical Z-stacks were acquired utilizing a Zeiss Axio Imager.M2 microscope equipped with an ApoTome.2 structured illumination module with the 20x objective and processed into maximum intensity projections.

### ACLMP organoids

ACLMP organoids^45^, containing salivary adenoid cystic carcinoma (ACC), colon epithelia, lung epithelia, mouse small intestine, and human pancreatic ductal adenocarcinoma (PDAC) organoid models, were manually segmented using coordinate bounding boxes from whole well displays to obtain single organoid level images. Precisely, a total of n = 80 organoids for ACC, n = 77 for colon epithelia, n = 176 for lung epithelia, n = 600 for mouse small intestine and finally n = 294 for PDACs. Images were equally split through an 80:20 train and test split. Models were tasked to separate organoids in a cross-sectional classification task into one of the five given categories. Source data used to segment and subsequently classify organoids are publicly available under https://osf.io/xmes4/.

### Colon organoids

Patient-derived colon organoids^44^ were generated using a Cytation five cell imaging multi-mode reader in the bright-field configuration with a 4x objective. Next, singular organoids were cropped using coordinate bounding boxes. The extracted images were subsequently organized into two distinct morphological classification tasks. For the opacity dataset, organoids were assessed based on luminal clearance. Organoids displaying a thin epithelium or clear lumen (transparent), and a thicker epithelium or lacking a clear lumen (opaque). Images were split into a training set (n = 1,981), validation set (n = 245), and test set (n = 252) using an 80:10:10 split. For the budding dataset, organoids were classified based on structural protrusions into budding (clear protrusion) and non-budding (mainly spherical) categories. This dataset was divided into a training set (n = 2,224), and a holdout test set (n = 252). All datasets were classified and split by the corresponding experts. The raw datasets can be accessed through the following repository at https://osf.io/42r3g/.

### Brain culture organoids

Brain organoids^42^, differentiated on differing culture devices (microfluidic device, orbital shaker, and static culture), were equally split into the three classes. In brief, 132 images were equally split on an 80:20 train and test ratio with n = 34 organoids for each category in the train and n = 8 organoids for each class in the test set. The prediction task was hence to truthfully separate organoids across their culture condition settings into one of the three categories. Brightfield images were obtained with a resolution of 2,464 × 2,056 pixels and subsequently resized to a resolution of 224 x 224 pixels for training of ViTAMIn-O. The original datasets used for this study can be obtained through: https://www.kaggle.com/datasets/burakkahveci/brain-organoid-and-embryoid-bodies-organolabeling.

### Data for performance evaluation

#### Brain organoid datasets

Two brain organoid datasets^63^ were collected containing normal-abnormal (72% normal, 28% abnormal) and normal-budding (60% normal, 40% budding) classes. The train and test data distribution was adopted with a split of n = 188 (train set) and n= 52 (test set) images for the first set in addition to n = 265 (train set) and n = 67 images for the second set. Labels were assessed by experts, Images were obtained in the brightfield configuration with a 10x objective and a resolution of 2,464 x 2,056 pixels using an inverted microscope equipped with a Zeiss Axiocam 705 colour camera. Datasets are accessible via the following links: https://www.kaggle.com/datasets/burakkahveci/brain-organoid-normal-and-abnormal-dataset and https://www.kaggle.com/datasets/burakkahveci/brain-organoid-budding-dataset.

#### PDLO organoids

Pancreatic duct-like organoids (PDLOs) were differentiated from human embryonic stem cells (hESCs) according to previously published protocols^64–67^. Daily live-cell brightfield imaging of whole well displays was performed up to day 11 of pancreatic progenitor differentiation into PDLOs using the Incucyte Live-Cell Analysis System. Differentiation outcomes on day 11 organoids were assessed based on overall morphological appearance either i) satisfactory or ii) unsatisfactory (Supplementary Fig. 3h). For training, labels were then retroactively assigned to the matched organoids on day 8 of differentiation at single organoid resolution. The overall task for Vitamin-O was then to predict satisfactory or unsatisfactory PDLO development, not at the endpoint but already at day 8 of differentiation. Overall, 200 organoids were tracked for each category, totaling in n = 400 organoid images, which were then equally split following 80:20 train and test fractions (n = 320 and n = 80 organoid images respectively). Imaging datasets can be obtained through the following link: https://doi.org/10.5281/zenodo.19466176.

#### Embryo datasets

The embryo developmental stage dataset^49^ encapsulates 5,500 timelapse images sourced from a clinical cohort and a publicly available repository. Images were obtained using an EmbryoScope time-lapse system under a 635 nm LED light source. The dataset is equally split across five developmental stages, namely 2-cell, 4-cell, 8-cell, morula, and blastocyst (totalling n = 1,100 images per class). To prevent data leakage, authors split the imaging data at the sequence level into a training set (n = 5,000 with 1,000 images per stage) and a holdout test set (with n = 500 and 100 images per stage). The blastocyst quality grading dataset^48^ encompasses n = 2,344 individual blastocyst images derived from n = 837 patients. Images were captured using an Olympus IX50 microscope at 400x magnification. To establish robust ground truth labels, an international consortium of expert embryologists annotated the images based on the clinical Gardner scoring system^68^. Consequently, the dataset is stratified into three classification tasks with expansion grade (EXP), inner cell mass (ICM), and trophectoderm (TE) quality. Datasets are openly accessible following the links: https://zenodo.org/records/14253170 and https://doi.org/10.6084/m9.figshare.20123153.v3.

### Automated concept-based explanation on organoids (OrgACE)

Concerning the proof of principle on automated concept discovery, a fine-tuning approach was adopted to enrich the latent embeddings of ViTAMIn-O. Next, the flattened patch embeddings across the otic dataset were clustered using k-means clustering (with k = 15, and n_init = 10). To present an intuitive visualization of these concepts, cluster assignments were inversely mapped to the original high resolution images. Relative patch grid coordinates were converted to absolute pixel bounding boxes, scaled by a customizable context expansion factor (> 1.0) to include surrounding morphological architecture and afterwards cropped. To quantify the biological enrichment of discovered concepts, image level annotations were assigned to their segmented patches. A contingency matrix of concept assignments versus biological classes was generated. To prevent majority class dominance, raw patch counts were first normalized by total class size, followed by a row wise percentage normalization to determine the precise underlying biological composition of each morphological concept.

## QUANTIFICATION AND STATISTICAL ANALYSIS

Demonstrating the differences in performance between deep learning models calls for a comprehensive comparison across a variety of metrics, especially in data scarce scenarios, which in addition, are subject to class imbalances. The following section dives into the differing metrics, which were employed in assessing the performance of ViTAMIn-O.

The per-category and overall accuracy describe the number of correct classifications over the total amount of ground-truth images and thus provides an intuitive ratio of correct to total predictions. It further serves as the measure by which the visualization of confusion matrixes can be performed.

The Area Under the Curve (AUC) score is a metric to evaluate a model’s performance concerning its discriminatory capability. The corresponding plot, the Receiver Operating Characteristic (ROC or more generally summed up as AUROC) under which the integral is calculated, illustrates varying threshold values concerning the true positive rate (sensitivity) against the true negative rate (specificity). Under this notion, an AUROC value of 0.5 represents chance, whilst values closer to 1 indicate clear separability capabilities of the evaluated model between positive and negative classes.

While the precision illustrates the ratio between TP (true positives) against the sum of TP and FP (false positives), the recall score is the ratio of the TP over the sum of TP and FN (false negatives).

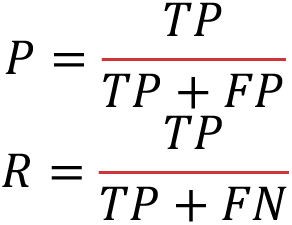

The F1 score is regarded as the harmonic mean between precision and recall with values between [0; 1], where 1 represents the best and 0 the worst possible score. For multiclass classification, the F1 score for all classes gets iteratively averaged or returned for all classes. It is calculated as follows:

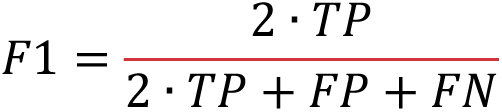

Matthew’s correlation coefficient (MCC), ranging from values in an interval between [-1; 1], encapsulates a comprehensive overview on the sensitivity, specificity, precision and the negative predictive value. A value closer to 1 corresponds to higher values in all four metrics, 0 represents chance and -1 an overall inverse prediction. The MCC score thus serves as a balanced measure on the quality of a model’s prediction across binary or multiclass classification tasks.

All statistical analyses were performed through independent two-sided independent t-test. P-values are described in the corresponding figures. We performed bootstrapping on the test sets, creating 1,000 resampled datasets to infer robust values of the 95% confidence intervals (CIs) of the metrics. AUROC scores and CIs were calculated iteratively on 1,000 instances, providing reliable estimates of the true predictive power of the model as well as its variability.

## ADDITIONAL RESOURCES

For more detailed descriptions on how to run ColabViTAMIn-O and deploy models from the Model hub as well as upload your customized models to the repository to share with researchers, visit the following tutorials. ColabViTAMIn-O tutorial: https://drive.google.com/file/d/1-brT1ZU8CdUeDluaQ0a8LqldHMeDwUgC/view?usp=sharing, and the ViTAMIn-O Model hub tutorial: https://drive.google.com/file/d/1P2VtXC22yHDKT0pjY3KcfcuIhdr5RP5M/view?usp=sharing. Figures 1, 3, and 5 were partially created with BioRender.com: Created in BioRender. Hamurcu, F. (2026) https://BioRender.com/ha8zqkq (Figure 1), Created in BioRender. Hamurcu, F. (2026) https://BioRender.com/09lkyxt (Figure 3A-C). Created in BioRender. Hamurcu, F. (2026) https://BioRender.com/dlg4dsf (Figure 5A).

