## Supplemental Information for "ViTAMIn-O: Democratizing computer vision-based machine learning for stem cell research"

**Document S1. Figures S1–S5**

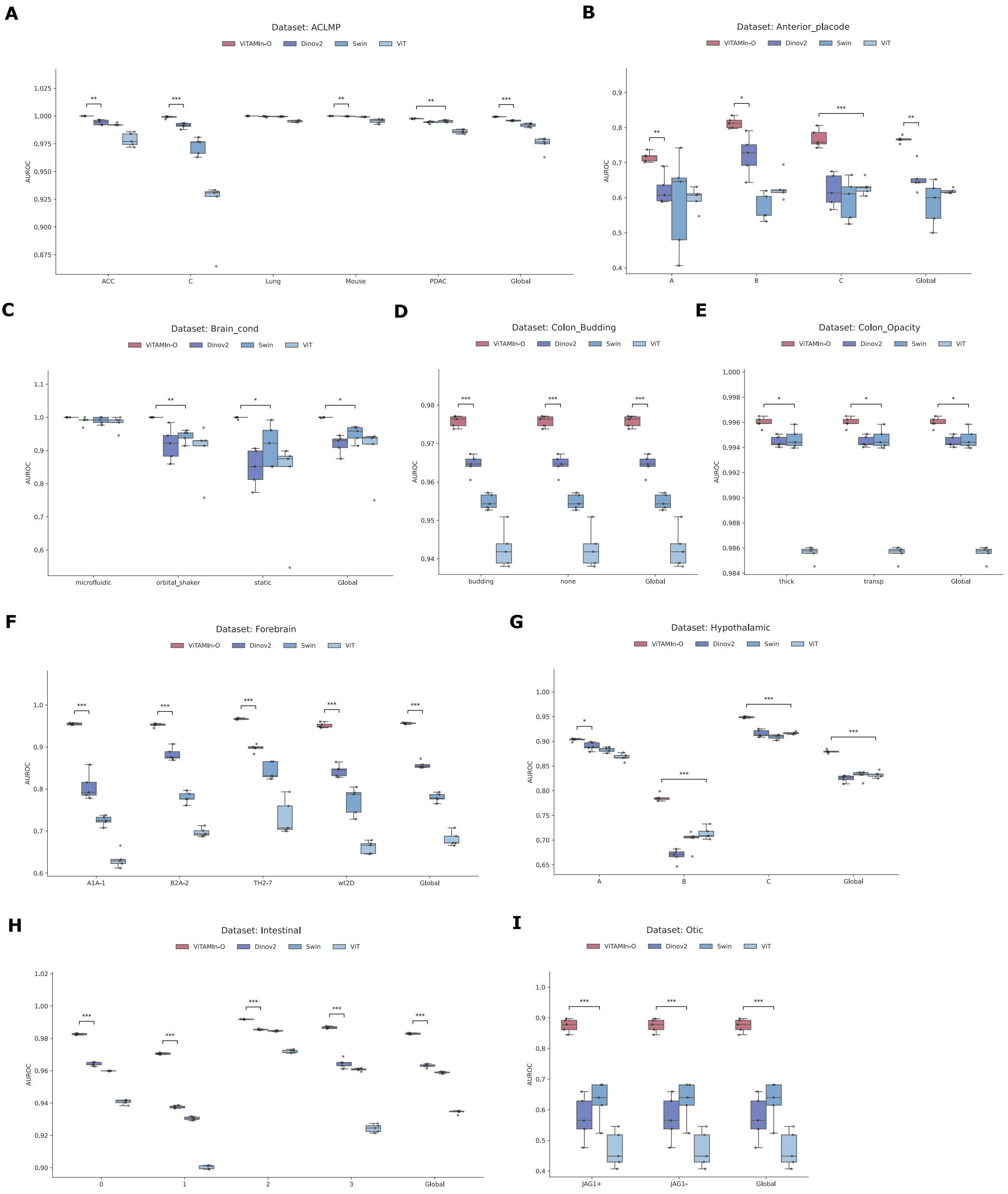
A
Dataset: ACLMP
ViTAMIn-O Dinov2 Swin ViT
AUROC
ACC
C
Lung
Mouse
PDAC
Global
B
Dataset: Anterior_placode
ViTAMIn-O Dinov2 Swin ViT
AUROC
A
B
C
Global
C
Dataset: Brain_cond
ViTAMIn-O Dinov2 Swin ViT
AUROC
microfluidic
orbital_shaker
static
Global
D
Dataset: Colon_Budding
ViTAMIn-O Dinov2 Swin ViT
AUROC
budding
none
Global
E
Dataset: Colon_Opacity
ViTAMIn-O Dinov2 Swin ViT
AUROC
thick
transp
Global
F
Dataset: Forebrain
ViTAMIn-O Dinov2 Swin ViT
AUROC
A1A-1
B2A-2
TH2-7
wt2D
Global
G
Dataset: Hypothalamic
ViTAMIn-O Dinov2 Swin ViT
AUROC
A
B
C
Global
H
Dataset: Intestinal
ViTAMIn-O Dinov2 Swin ViT
AUROC
0
1
2
3
Global
I
Dataset: Otic
ViTAMIn-O Dinov2 Swin ViT
AUROC
JAG1+
JAG1-
Global

**S1 Fig. Per-class AUROC metrics on hold out test dataset under the linear probing configuration.** (A-I) Global Area Under the Curve Receiver Operating Characteristic (AUROC) comparisons of ViTAMIn-O against, DINOv2-large, Swin-large, or ViT-large across all nine and 28 per-class organoid prediction tasks. (A) ACLMP ViTAMIn-O vs. DINOv2: t = 16.0797, p = 6.6460e-06. (B) Anterior placode ViTAMIn-O vs. DINOv2: t = 6.2946, p = 2.1880e-03. (C) Brain culture conditions ViTAMIn-O vs. Swin: t = 4.5840, p = 1.0042e-02. (D) Colon Budding ViTAMIn-O vs. DINOv2: t = 8.6000, p = 9.3168e-05. (E) Colon Opacity ViTAMIn-O vs. Swin: t = 3.3250, p = 1.5363e-02. (F) Forebrain ViTAMIn-O vs. DINOv2: t = 24.7044, p = 8.2683e-06. (G) Hypothalamic-pituitary ViTAMIn-O vs. ViT: t = 14.3615, p = 8.3330e-06. (H) Intestinal ViTAMIn-O vs. DINOv2: t = 38.0028, p = 1.6206e-07. (I) Otic ViTAMIn-O vs. Swin: t = 8.0353, p = 5.3876e-04. Data entries are presented as mean ± standard deviation across five independent runs. Across all runs, we selected a batch size of 32 (batch_size = 32) and epoch size of 20 (epochs = 20). Statistical analyses were carried out using Welch's t-test.

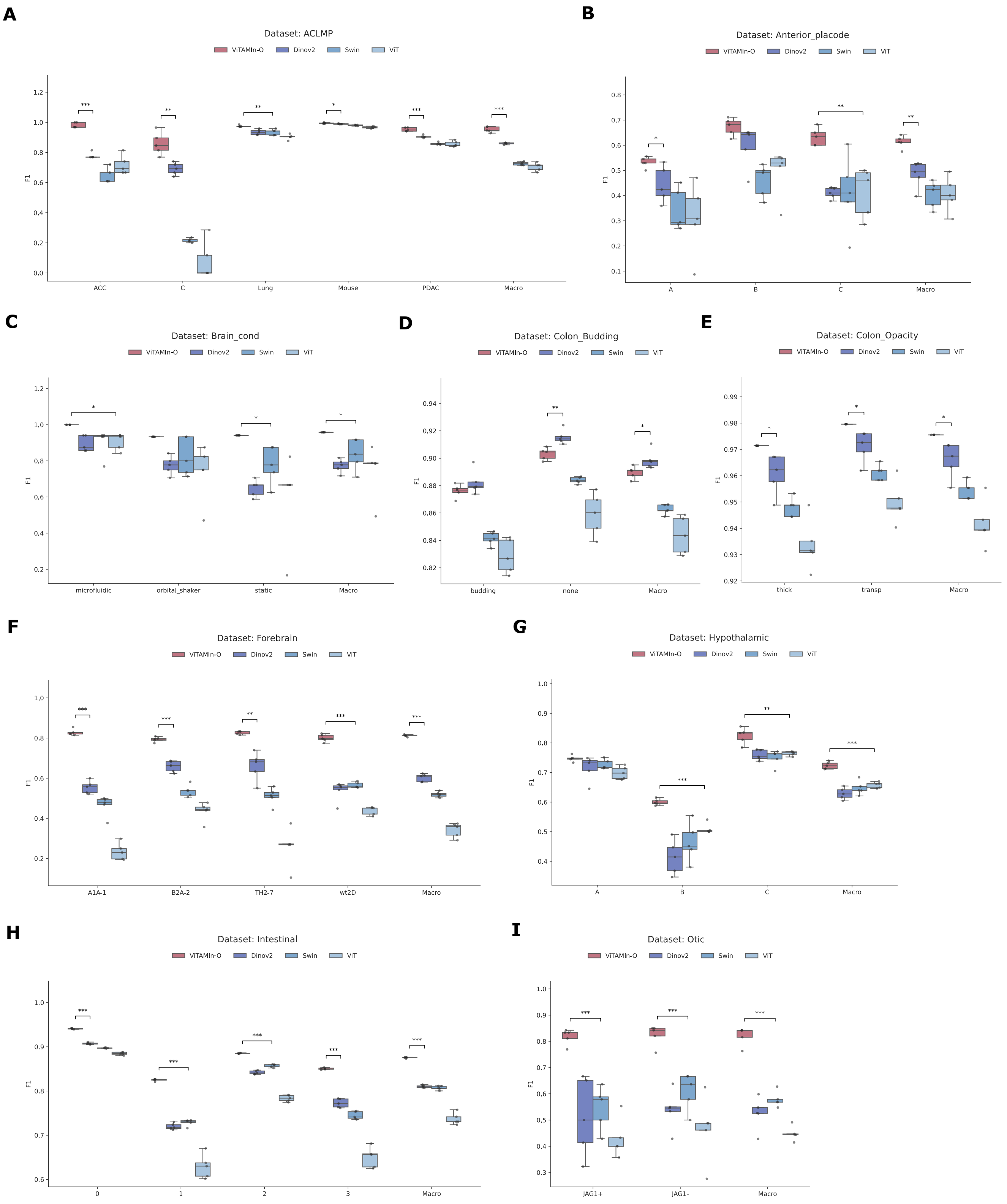
A
Dataset: ACLMP
ViTAMIn-O Dinov2 Swin ViT
F1
ACC C Lung Mouse PDAC Macro
B
Dataset: Anterior_placode
ViTAMIn-O Dinov2 Swin ViT
F1
A B C Macro
C
Dataset: Brain_cond
ViTAMIn-O Dinov2 Swin ViT
F1
microfluidic orbital_shaker static Macro
D
Dataset: Colon_Budding
ViTAMIn-O Dinov2 Swin ViT
F1
budding none Macro
E
Dataset: Colon_Opacity
ViTAMIn-O Dinov2 Swin ViT
F1
thick transp Macro
F
Dataset: Forebrain
ViTAMIn-O Dinov2 Swin ViT
F1
A1A-1 B2A-2 TH2-7 wt2D Macro
G
Dataset: Hypothalamic
ViTAMIn-O Dinov2 Swin ViT
F1
A B C Macro
H
Dataset: Intestinal
ViTAMIn-O Dinov2 Swin ViT
F1
0 1 2 3 Macro
I
Dataset: Otic
ViTAMIn-O Dinov2 Swin ViT
F1
JAG1+ JAG1- Macro

**S2 Fig. Per-class F1 score metrics on hold out test dataset under the linear probing configuration.** (A-I) Global F1 score comparisons of ViTAMIn-O against, DINOv2-large, Swin-large, or ViT-large across all nine and 28 per-class organoid prediction tasks. (A) ACLMP ViTAMIn-O vs. DINOv2: $t = 10.5630$, $p = 1.6959e-04$. (B) Anterior placode ViTAMIn-O vs. DINOv2: $t = 4.9636$, $p = 3.0549e-03$. (C) Brain culture conditions ViTAMIn-O vs. Swin: $t = 3.1560$, $p = 3.4314e-02$. (D) Colon Budding DINOv2 vs. ViTAMIn-O: $t = 2.5019$, $p = 4.1523e-02$. (E) Colon Opacity ViTAMIn-O vs. DINOv2: $t = 3.1922$, $p = 3.3147e-02$. (F) Forebrain ViTAMIn-O vs. DINOv2: $t = 23.3107$, $p = 6.1388e-06$. (G) Hypothalamic-pituitary ViTAMIn-O vs. ViT: $t = 9.3327$, $p = 1.7319e-05$. (H) Intestinal ViTAMIn-O vs. DINOv2: $t = 45.1014$, $p = 5.7589e-08$. (I) Otic ViTAMIn-O vs. Swin: $t = 11.9484$, $p = 2.5879e-06$. Data entries are presented as mean ± standard deviation across five independent runs. Across all runs, we selected a batch size of 32 (batch_size = 32) and epoch size of 20 (epochs = 20). Statistical analyses were carried out using Welch's t-test.

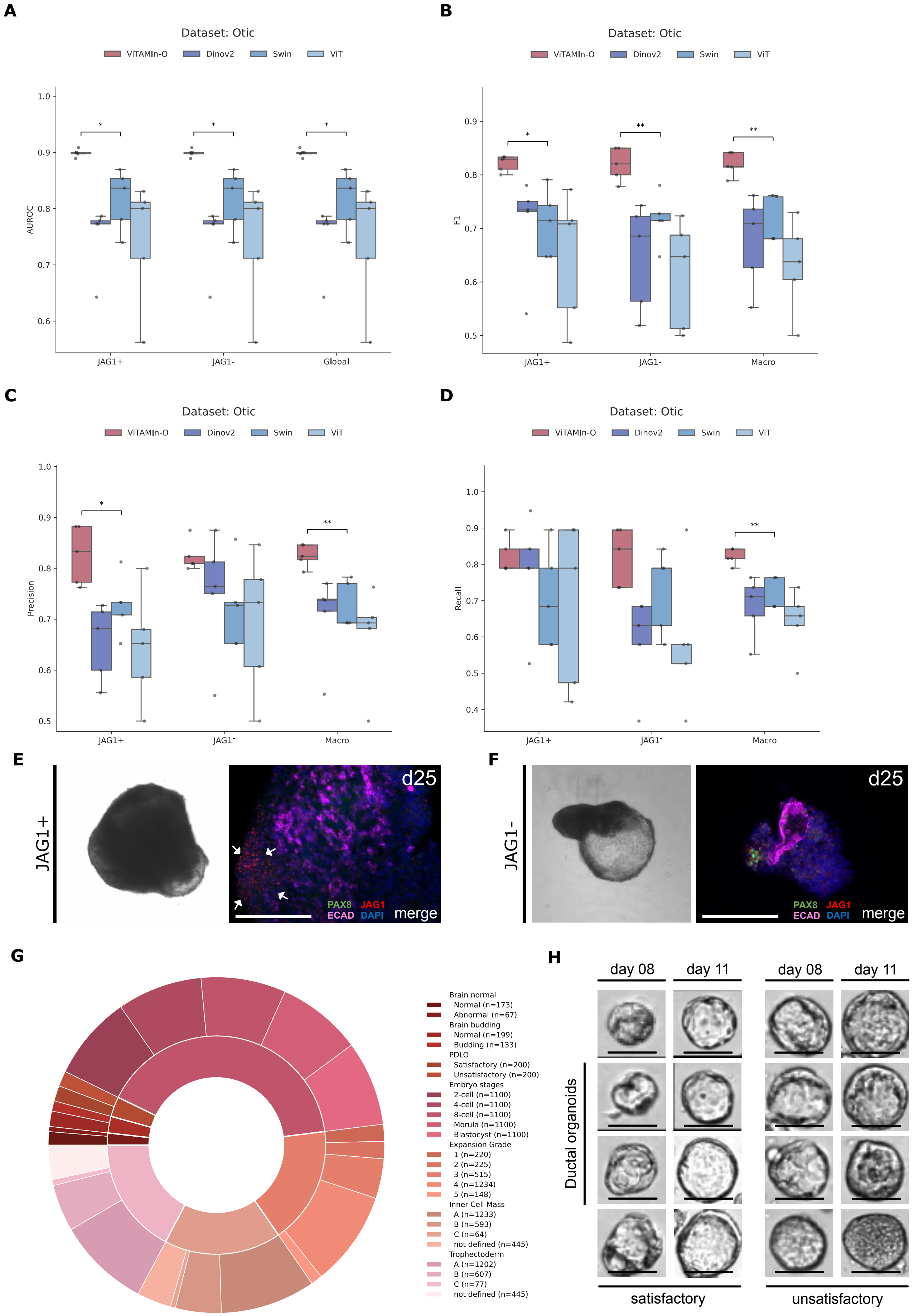
A
Dataset: Otic
ViTAMIn-O Dinov2 Swin ViT
AUROC
JAG1+
JAG1-
Global
B
Dataset: Otic
ViTAMIn-O Dinov2 Swin ViT
F1
JAG1+
JAG1-
Macro
C
Dataset: Otic
ViTAMIn-O Dinov2 Swin ViT
Precision
JAG1+
JAG1-
Macro
D
Dataset: Otic
ViTAMIn-O Dinov2 Swin ViT
Recall
JAG1+
JAG1-
Macro
E
JAG1+
d25
PAX8 JAG1
ECAD DAPI merge
F
JAG1-
d25
PAX8 JAG1
ECAD DAPI merge
G
Brain normal
Normal (n=173)
Abnormal (n=67)
Brain budding
Normal (n=199)
Budding (n=133)
PDLO
Satisfactory (n=200)
Unsatisfactory (n=200)
Embryo stages
2-cell (n=1100)
4-cell (n=1100)
8-cell (n=1100)
Morula (n=1100)
Blastocyst (n=1100)
Expansion Grade
1 (n=220)
2 (n=225)
3 (n=515)
4 (n=1234)
5 (n=148)
Inner Cell Mass
A (n=1233)
B (n=593)
C (n=64)
not defined (n=445)
Trophectoderm
A (n=1202)
B (n=607)
C (n=77)
not defined (n=445)
H
day 08
day 11
day 08
day 11
Ductal organoids
satisfactory
unsatisfactory

**S3 Fig. Per-class metrics on the otic dataset under the fine-tuning configuration and visualization of PDLO organoid differentiation prediction paradigm.** (A-D) Global AUROC, F1, Precision and Recall metrics on the hold-out test set of otic organoids. Models were finetuned over the course of 20 epochs (epochs = 20) with a batch size of 32 (batch_size = 32). Data entries are presented as mean ± standard deviation across five independent runs and statistical analyses were performed using Welch's t-test. (A) AUROC ViTAMIn-O vs. Swin: $t = 3.4008$, $p = 2.5917e-02$. (B) F1 score ViTAMIn-O vs. Swin: $t = 4.9356$, $p = 2.7106e-03$. (C) Precision ViTAMIn-O vs. Swin: $t = 4.3242$, $p = 5.4021e-03$. (D) Recall ViTAMIn-O vs. Swin: $t = 4.8507$, $p = 2.9248e-03$. (E-F) Exemplary matched PDLO organoids on day 8 and day 11 of differentiation, grouped after endpoint differentiation outcome results. (E) Satisfactory organoid with translucent edges and denser cores. (F) Unsatisfactory organoids with an imposing translucent cyst-like structure. Scale bars 100 µm. (G) Simplified schematic of ViTAMIn-O's overall testing data structure. (H) Single PDLO organoids are tracked from day 8 onwards and retrospectively assigned labels corresponding to their differentiation endpoint morphology on day 11. ViTAMIn-O is trained to predcit the overall outcome based on snapshot images of earlier timepoints during the course of differentiation.

c

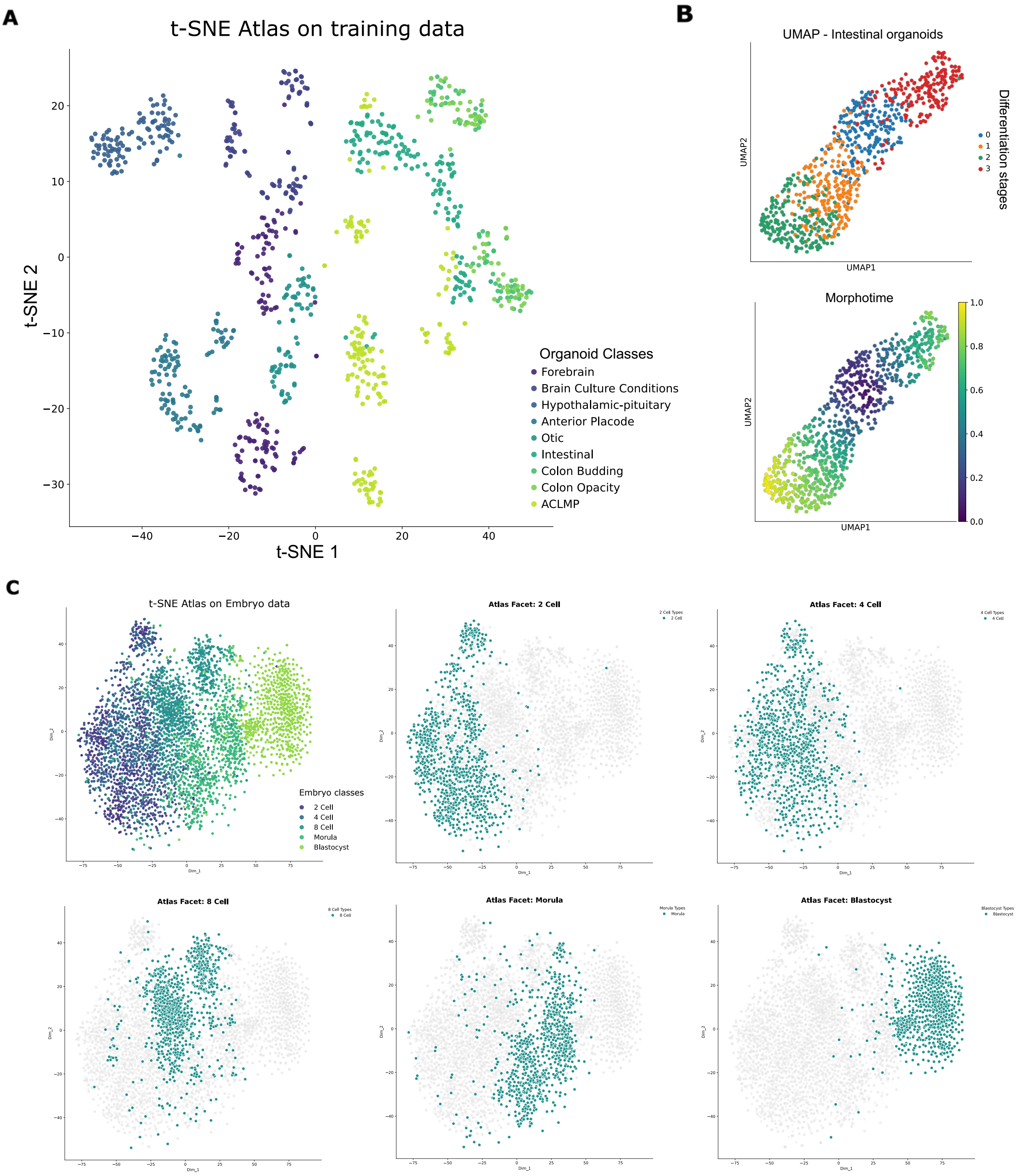
A
t-SNE Atlas on training data
t-SNE 2
t-SNE 1
Organoid Classes
Forebrain
Brain Culture Conditions
Hypothalamic-pituitary
Anterior Placode
Otic
Intestinal
Colon Budding
Colon Opacity
ACLMP
B
UMAP - Intestinal organoids
UMAP2
UMAP1
Differentiation stages
0
1
2
3
Morphotime
C
t-SNE Atlas on Embryo data
Embryo classes
2 Cell
4 Cell
8 Cell
Morula
Blastocyst
Atlas Facet: 2 Cell
Atlas Facet: 4 Cell
Atlas Facet: 8 Cell
Atlas Facet: Morula
Atlas Facet: Blastocyst

**S4 Fig. Dimensionality reduction on various imaging sets.** (A) Global t-SNE projection of ViTAMIn-O's feature embeddings encompassing all nine organoid datasets, encountered during pretraining. The number of individual points is normalized across all categories to balance the quantitative heterogeneity. (B) UMAP and Morphotime analysis on the isolated intestinal organoid dataset. Baseline spheroids (blue, category 0), diverge towards either a healthy developmental vector of early (orange, category 1) and late organoids (green, categories 2) or towards the cystic morphological type (red, category 3). Morphotime analysis further underscores the morphologically grounded trajectorial distance between late and cystic organoids. (C) t-SNE map on sequential embryo imaging data from the 2-cell to blastocyst stage. ViTAMIn-O's embeddings naturally mapped the intermediate steps of differentiation without pretraining or fine-tuning. All hyperparameters needed to derive the presented maps were generated using 20 nearest neighbor (n_neighbors = 20), a minimum distance of 0.1 (min_dist = 0.1), and the cosine distance metric for UMAP. For t-SNE, a perplexity of 30 (perplexity = 30) as well as 1.000 iterations (iterations = 1000) were use.

| External dataset | Metric | Albumentations pipeline | Standard ViTAMIn-O pipeline |
|---|---|---|---|
| Brain Budding Organoids | AUROC | 0.987 ± 0.004 | **0.991** ± **0.003** |
| Brain Normal Organoids | AUROC | 0.976 ± 0.010 | **0.999** ± **0.001** |
| PDLO Organoids | AUROC | 0.991 ± 0.003 | **0.993** ± **0.002** |

**S5 Fig. Data augmentation comparisons between Albumentations and the ViTAMIn-O pipeline.** Our data augmentation algorithm, incorporated into the ColabViTAMIn-O platform and used during pretraining and downstream analysis, constantly outperforms the Albumentations strategy on the brain budding, brain anormal and PDLO organoid datasets. ViTAMIn-O was trained in the linear probing configuration over the course of 20 epochs (epochs = 20) with a batch size of 32 (batch_size = 32). Displayed results are the means ± standard deviation across five independent runs.
